# Epigenetic inheritance of gene-silencing is maintained by a self-tuning mechanism based on resource competition

**DOI:** 10.1101/2022.06.15.496123

**Authors:** Omer Karin, Eric A. Miska, Benjamin D. Simons

## Abstract

Biological systems can maintain memories over long timescales, with examples including memories in the brain and immune system. It is currently unknown how functional properties of memory systems, such as memory persistence, can be established by biological circuits. To address this question, we focus on transgenerational epigenetic inheritance in C. elegans. In response to a trigger, worms silence a target gene for multiple generations, resisting strong dilution due to growth and reproduction. Silencing may also be maintained indefinitely upon selection according to silencing levels. We show that these properties imply fine-tuning of biochemical rates in which the silencing system is positioned near the transition to bistability. We demonstrate that this behavior emerges from a generic mechanism based on competition for synthesis resources, which leads to self-organization around a critical state with broad silencing timescales. The theory makes distinct predictions and offers insights into the design principles of long-term memory systems.

## Introduction

The ability to record and maintain memories is one of the distinctive and most remarkable features of living systems. Examples include the ability of the brain to record past experiences, the immune system to remember pathogen encounters, and many organisms to transfer epigenetic information across generations (Ashe et al., 2012; Houri-Zeevi and Rechavi, 2017; Iwasaki and Paszkowski, 2014; Kandel, 2001; Macallan et al., 2017; Rechavi and Lev, 2017).

From the perspective of systems biology, it is important to understand how the functional properties of long-term memory systems can be implemented by biological interactions, such as networks of biochemical reactions. This challenge is exemplified by transgenerational transcriptional silencing in *C. elegans* worms. When exposed to a double-stranded RNA (dsRNA) trigger, worms will silence its homologous gene. Remarkably, when expressed in the germline, this silencing can persist over multiple generations even in the absence of the original trigger (Fire et al., 1998). Transgenerational silencing also occurs in response to various endogenous triggers (Ashe et al., 2012; Beltran et al., 2020; Lev et al., 2019; Luteijn et al., 2012; Rechavi et al., 2014). At the molecular level, there are two processes that contribute to silencing – the production and amplification of 22-nucleotide-long short-interfering RNAs (22G siRNAs) and the deposition of silencing histone modifications including the heterochromatin mark H3K9me3 (Rechavi and Lev, 2017; Woodhouse and Ashe, 2020) (Figure 1A,B). Transgenerational transcriptional silencing allows *C. elegans* to protect itself from foreign agents such as viruses (Rechavi et al., 2011) and transposable elements (Bagijn et al., 2012), and to transmit environmental information across generations (Klosin et al., 2017; Rechavi et al., 2014). The relatively short generation time of *C. elegans,* together with the ability to control memory induction, makes it a particularly attractive model to study memory at both the molecular (Frolows and Ashe, 2021) and phenomenological (Beltran et al., 2020; Houri-Zeevi et al., 2020) level.

**Figure 1.**
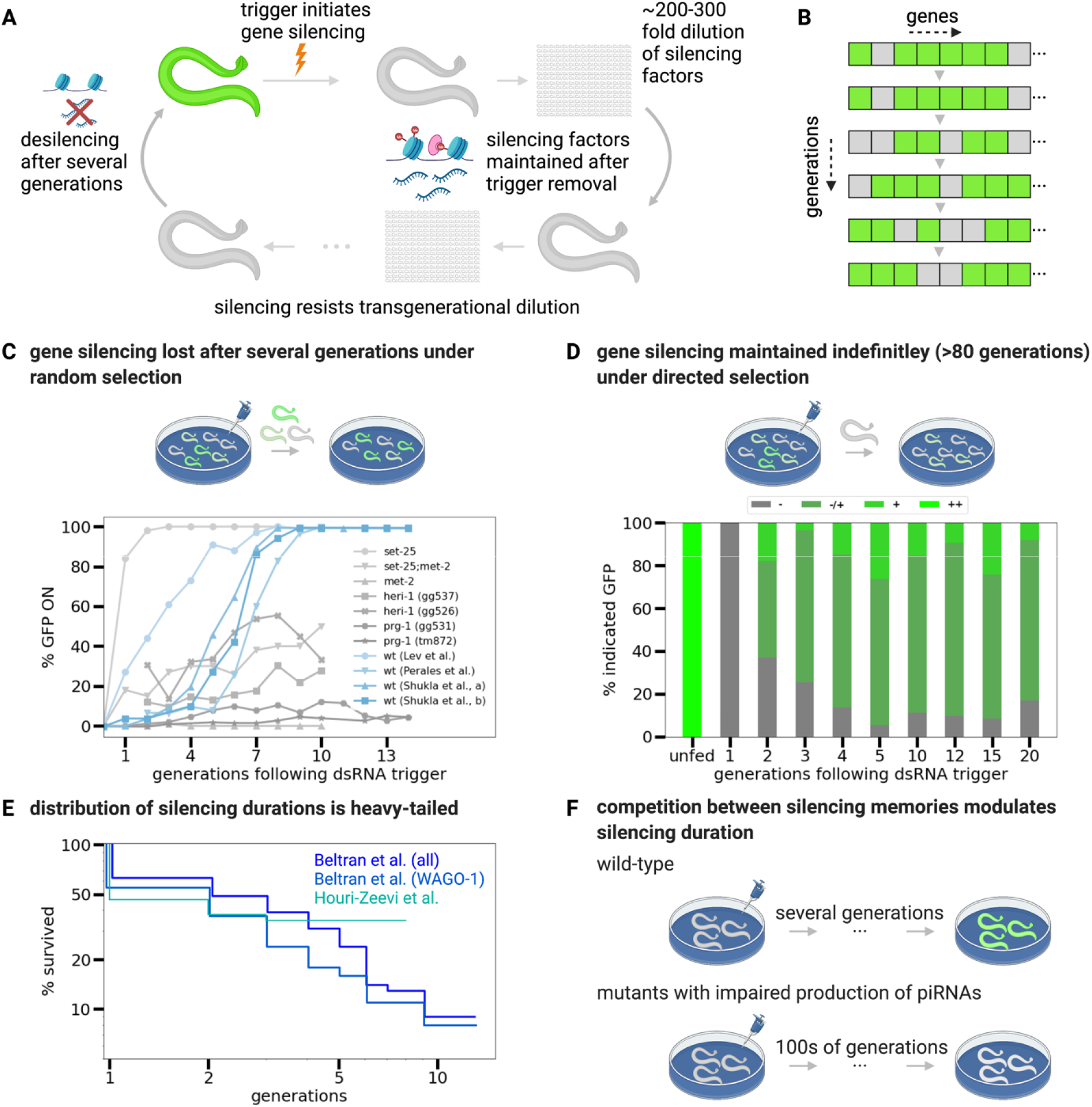
Experimental observations of transgenerational gene silencing in *C. elegans* constrain possible response mechanisms. (A) Transgenerational co-transcriptional silencing of *gfp* expression in *C. elegans* is initiated following the presentation of a dsRNA trigger and can be tracked at the level of individual animals. Silencing is associated with production, transmission, and maintenance of siRNAs and silencing chromatin marks (H3K9me3) that persists over multiple generations, resisting many-fold dilution due to reproduction of worms. Silencing terminates following the loss of the relevant siRNA molecules and silencing chromatin modifications. (B) Schematic illustrating that stochastic silencing (depicted as a transition from color gray) also occurs in many endogenous genes (depicted as distinct colors) and may persist over the timescale of multiple generations. (C) Percentage of worms that show *gfp* expression as a function of generation number following induced silencing. In the wild type (wt), silencing persists for around generations, when averaged over the population of worms. (Blue lines denote wt strains, data from (Lev et al., 2017; Perales et al., 2018; Shukla et al., 2021); note that Shukla et al., a,b refer to measurements on wt strains from Figure S1A,B). However, some mutant strains show altered average silencing times that can be shorter, such as the *set-25* knockdown where *T* ≈ 1 generation, or longer, such as the *met-2* knockdown where *T* ≳ 30 generations. (Gray lines denote mutant strains, data from (Lev et al., 2017; Perales et al., 2018; Shukla et al., 2021)). (D) Although the average persistence time of *gfp* silencing across generations is limited, if worms with the highest degree of silencing are continuously selected from the ensemble at each generation, silencing levels of their progenies converge towards a broad and stationary distribution that can be maintained indefinitely (panel adapted from Figure 1 of (Vastenhouw et al., 2006); ++ corresponds to strong *gfp* signal, while +, +/−, and – correspond to increasingly weaker *gfp* signal). (E) Distribution of silencing durations for a given lineage is “heavy-tailed”, showing a slow (power-law like) decay at long silencing times (data on *gfp* silencing in individual lineages from (Houri-Zeevi et al., 2020) (turquoise), data on siRNA epimutation duration from (Beltran et al., 2020) (shades of blue). (F) Amplification and maintenance of silencing machinery (e.g., siRNAs) is limited by shared synthesis components. Resource competition is tightly associated with silencing duration, and mutants where competition between silencing memories is modulated (e.g., animals that lack piRNAs) have much longer silencing durations.

Here, we make use of the known properties of this system to address two basic and generic questions: (1) How can a biological memory system establish and tune a prolonged time of memory duration? (2) Are there features of the memory system that are beneficial for retaining relevant memories? Both questions are functionally relevant for the epigenetic silencing system. Silencing in wild-type worms is retained for *T* ≈ 4 – 7 generations following the removal of the trigger, which is much longer than the typical timescale of dilution of the silencing factors by growth and reproduction (Houri-Ze’evi et al., 2016), yet much shorter than the silencing durations observed in certain mutants (Lev et al., 2017; Shukla et al., 2021). Tuning the timescale *T* allows the system to balance the potential benefits of long-term silencing with the aberrant accumulation of possibly irrelevant silencing memories. Additionally, there is variation in silencing levels between individual worms, and experimental selection on this variation allows silencing to be retained indefinitely in the population (Vastenhouw et al., 2006), implying that the variation is heritable. It is unclear how these features (tuning of memory duration over prolonged timescales, and heritable variation that allows for selection) can be implemented within the framework of a biochemical network.

Here, we show that the functional features of the transgenerational gene silencing memory system in *C. elegans* are attained through a competition mechanism that maintains the dynamics of silencing memories near a transition to bistability. To develop this framework, we first consider a minimal model based on dynamical experimental measurements, where 22G-siRNA amplification is governed by a bistable switch, and there is negative feedback due to the deposition of silencing chromatin marks. By itself, this circuit can behave either as a bistable switch (silencing factors are on/off) or provide a monostable decaying response (once activated, silencing factors decay towards a stable off state). However, silencing memories are not maintained in isolation, but rather share various synthesis components such as RNA-dependent RNA polymerases (RdRP) and Argonaute (AGO) proteins. Competition for these resources engages the collective behavior of many silencing memories, where their accumulation reduces the available amplification capacity. We show that, when accounting for such competition, the dynamics of silencing memories becomes self-organized in a narrow range near the transition to bistability in a manner that provides a tunable delay time, whose magnitude may be much longer than the timescale of the turnover of the underlying molecular components. Within this framework, the delay time is robust to noise and fluctuations in circuit parameters. Self-organization near the transition to bistability results in large, static, variation of gene silencing levels between individual worms, allowing for selection according to silencing levels. The memory model provides a unifying and predictive explanation of multiple – often paradoxical – observations on the transcriptional silencing of exogenous and endogenous targets in *C. elegans*. Based on the simplicity of the model assumptions, we postulate that this mechanism of *self-tuned criticality by competition* may prevail in other memory systems. We therefore conclude by outlining the general hallmarks and predictions of the model.

## Results

### A model for transgenerational epigenetic inheritance based on interactions between siRNAs and silencing chromatin marks

A well-established system for studying transgenerational inheritance involves the silencing of a constitutively expressed *gfp* reporter gene (Ashe et al., 2012; Buckley et al., 2012; Burkhart et al., 2011; Houri-Ze’evi et al., 2016; Houri-Zeevi et al., 2020; Luteijn et al., 2012; Vastenhouw et al., 2006). Silencing of a *gfp* reporter provides a quantitative proxy for the degree of silencing in an individual animal, and how silencing evolves in a population over generations. This approach has been used extensively to explore the influence on inheritance dynamics of selection (Houri-Zeevi et al., 2020; Vastenhouw et al., 2006; Woodhouse et al., 2018) and genetic background (Burton et al., 2011; Lev et al., 2017; Perales et al., 2018; Shirayama et al., 2012; Shukla et al., 2021; Spracklin et al., 2017). *gfp* silencing experiments show that, following its induction, silencing is typically reversed in wild-type animals within around *T* ≈ 3 – 7 generations (Figure 1C). Moreover, mutant strains can show altered inheritance patterns, including long-term silencing that can persist for tens of generations or more (Figure 1C).

Most transgenerational silencing experiments (including those reproduced in Figure 1C) involve the transfer of random progeny to new plates at each generation, with the population of worms at each generation showing various degrees of silencing. However, Vastenhouw et al. found that, by selecting worms that show the highest degree of silencing at each generation, a stable inheritance pattern of silencing is established among progenies that can last for at least 80 generations (Vastenhouw et al., 2006) (Figure 1D). Notably, in every generation, the same continuous distribution of silencing states is replicated or re-established. These findings suggest that the silencing of a target gene that provides a selective benefit to an organism can be maintained for much longer than the average *T* ≈ 3 – 7 generation number for the population as a whole.

Further quantitative insight into transgenerational inheritance dynamics comes from a recent study by Houri-Zeevi et al., where inheritance was tracked in multiple lineages composed of more than 20,000 worms (Houri-Zeevi et al., 2020). Their study revealed that, in contrast to the variability in silencing levels observed at the population level by Vastenhouw et al. under selection, the process of de-silencing occurs approximately uniformly in the descendants of each worm. They also showed that some individual lineages retain silencing for much longer than the average duration time *T* of the population, as indicated by the distribution of silencing times that show a “heavy-tail” marked by a power-law like dependence (Figure 1E). Indeed, a similar heavy-tailed distribution has been observed in the duration of silencing epimutations in endogenous genes, which occur following stochastic initiation triggers (Figure 1E).

Silencing duration times also appear to be modulated by resource competition. The amplification of siRNAs requires the synthesis of resources such as AGO proteins and RdRP complexes. As these are shared between many small RNA molecules, they limit the overall synthesis rate, leading to competition for resources. Evidence for such competition is extensive in *C. elegans* (Duchaine et al., 2006; Houri-Ze’evi et al., 2016; Lee et al., 2006; Pavelec et al., 2009; Sarkies et al., 2013; Shirayama et al., 2012; Zhuang and Hunter, 2012), and experiments where long-lasting inheritance of silencing was demonstrated were accompanied by large-scale global changes in siRNAs (Lev et al., 2017; Shukla et al., 2021). Specifically, a recent study demonstrated that, in the absence of piRNAs (a major source of endogenous small RNAs that compete with siRNAs over shared synthesis resources), silencing can persist for at least hundreds of generations (Figure 1F) (Shukla et al., 2021).

Taken together, these experimental findings (summarized in Figure 1) represent the major hallmarks of gene silencing dynamics in *C. elegans* and constitute a set of exacting and diverse constraints on the underlying mechanisms that regulate this silencing memory system. Here, we focused on using these observations to identify how the functional properties of the epigenetic memory system may arise from interactions of the underlying biochemical components. Specifically, we placed emphasis on the two core molecular components of transgenerational inheritance: the concentration of the target-specific secondary siRNA pool (22G siRNAs) and the concentration of H3K9me3 on the target gene, both of which can be measured using small RNA sequencing (sRNAseq) and chromatin immunoprecipitation followed by sequencing (ChIPseq), respectively (Figure 2). We considered experiments where both quantities were measured in populations of worms following a dsRNA trigger (see Figure 2A,B for details) (Gu et al., 2012; Lev et al., 2017). Following exposure of wild-type animals to the dsRNA trigger, the P0 generation, the concentration of the target-specific siRNA pool (denoted *g*) and concentration of H3K9me3 around the target gene (denoted *h*), in populations of whole animals, follow a specific trajectory in phase-space (Figure 2A,B): An initial jump in *g*, followed by an increase in *h*, establishes a high level of gene silencing in P0. In subsequent generations, *g* decays slowly, while complete de-silencing in later generations is accompanied by reversal of both *g* and *h* back to their baseline (pre-trigger) values, closing the trajectory. In the *met-2* knockout animals, which show stable silencing of *gfp* over many generations, high levels of *g* and *h* are maintained at an effective transgenerational “stable fixed point” (Figure 2C).

**Figure 2.**
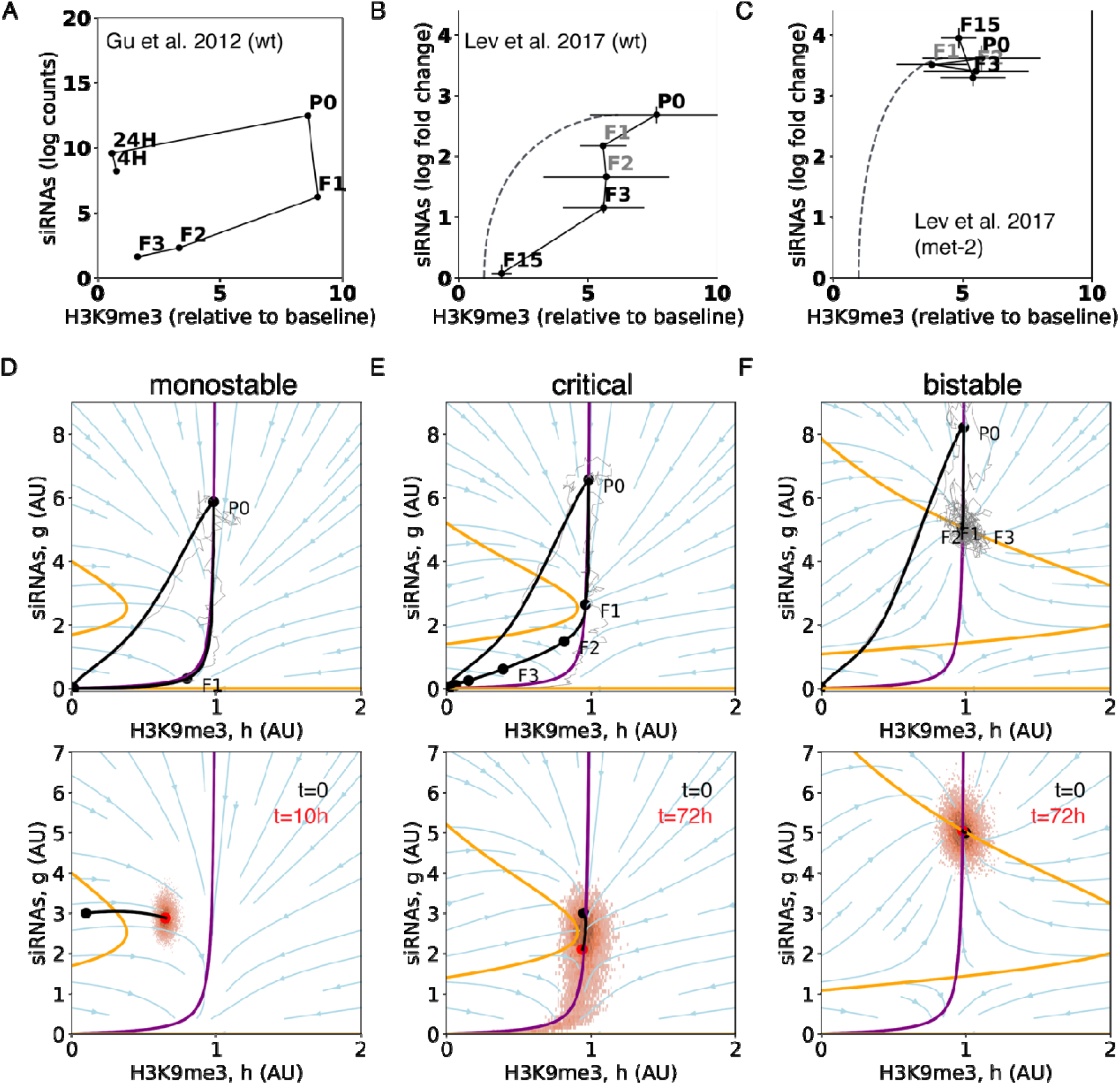
*Toggle-Inhibitor (TI)* model captures transgenerational dynamics of gene silencing factors. (A-C) Measured transgenerational dynamics of target-specific siRNAs and the accumulation of H3K9me3 marks on the target gene: in (A) wild-type (*wt*) worms following *lin-15B* dsRNA trigger (data from (Gu et al., 2012)); and (B) in *wt* worms and (C) *met-2* worms following *gfp* dsRNA trigger (data from (Lev et al., 2017)). Note that siRNA data was not available for F1 and F2 in panels B and C, and so was imputed using measured P0 and F3 siRNA levels. Measurements of H3K9me3 were performed using ChIP assays, while siRNA counts were quantified using siRNA-seq (Methods, see (Gu et al., 2012; Lev et al., 2017) for details). Error bars denote SEM. (D-F) Phase plane representations of the noise-free *TI* model in the three regimes: (D) monostable, (E) near saddle-node bifurcation or critical, and (F) bistable. Nullclines for and are shown as lines, parameterized by – — (purple) and – — (orange), respectively. The model has a stable unsilenced state at the origin while, for appropriate parameters, a silenced stable state (at high) can emerge. The upper panels depict the locus of trajectories of and obtained from stochastic simulations of the *TI* model with noise averaged over the population (black lines), as well as an example of an individual worm lineage (thin gray line). The lower panels depict the distribution of end states after one generation or less, starting from a single ancestral state positioned at the given point in phase space (black dot), representing the evolution over time of the silencing state (red dot represents the center-of-mass of the distribution of end states). Note that, in the monostable regime, the coordinates of all progenies are remote from the parent, moving towards the de-silencing state. Simulation parameters are provided in *Table 1.*

Altogether, the observed dynamics are consistent with a model of siRNAs and H3K9me3 with the following minimal set of processes (Figure 2D-F): (i) siRNAs amplify cooperatively (Groenenboom et al., 2005; Houri-Zeevi et al., 2020), resulting in bistability at low levels of H3K9me3; (ii) siRNAs promote the placement of H3K9me3 silencing marks (Gu et al., 2012; Holoch and Moazed, 2015); (iii) H3K9me3 modifications induce inhibition of the siRNA pool, e.g., due to transcriptional inhibition, which depletes the template mRNA, or due to the recruitment of allosteric inhibitors (Perales et al., 2018); and (iv) silencing factors decay on the transgenerational timescale by growth and reproduction. While it is possible to envisage additional mechanisms and processes acting on siRNAs and silencing marks, as well as other factors contributing to silencing, we propose that (i)-(iv) constitute the core set of processes that both capture the dynamics of siRNAs and H3K9me3 and, as we will show below, the observed transgenerational silencing features presented in Figure 1.

Mathematically, based on processes (i)-(iv), we propose that the dynamics of the siRNA pool and H3K9me3 concentration can be captured by a minimal model based on two coupled noisy rate equations:

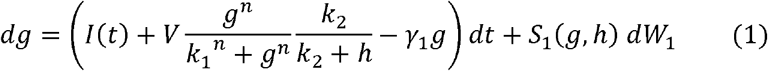

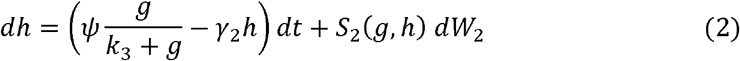

Here, *I*(*t*) denotes a time-dependent pulse-like induction term representing the initial trigger arising from the ingestion of bacteria expressing dsRNA in P0. The second production term, describing the cooperative amplification of 22G siRNA, scales in proportion to 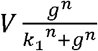, having a maximum amplification capacity *V* (that may be limited for example by available synthesis machinery), cooperativity *n*, and a half-maximal amplification activity at *g* = *k*_1_. The amplification term is modulated by an inhibitory factor, 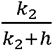, associated with silencing chromatin marks, with half-maximal strength at *h* = *k*_2_. 22G siRNAs stimulate the production of silencing chromatin marks with a maximum synthesis capacit *ψ* and halfway activity *g* = *k*_3_. We note that, on the transgenerational timescale, the production terms effectively combine both the production of silencing factors and their loading onto oocytes. Finally, *γ*_1_*γ*_2_ denote the respective decay rates of *g*, *h* due to growth and reproduction. In addition to these elements that act deterministically on the dynamics of *g*, *h*, to account for stochastic fluctuations between individuals due to variations in growth rate, as well as fluctuations in the expression of relevant factors such as heat shock proteins (Houri-Zeevi et al., 2020), we have included noise terms *S*_1_(*g*, *h*)*dW*_1_, *S*_2_(*g*, *h*)*dW*_2_, where *W*_1_, *W*_2_ are Wiener processes (Van Kampen, 1992). (Note that, for the stochastic simulations of the model, we consider multiplicative noise, setting *S*_1_(*g*, *h*) = *σ*_1_*g*., *S*_2_(*g*, *h*) = *σ*_2_*h*, with *σ*_1_, *σ*_2_ characterizing the strength of the noise.) Together, these coupled rate equations comprise the *Toggle-Inhibitor (TI)* model.

### Experimental observations on transgenerational dynamics are explained by a fine-tuned toggle-inhibitor model

The *TI* model (Eqs. 1,2) constitutes an excitable circuit that combines positive and negative feedback, a motif prevalent in many biological systems (Rué and Garcia-Ojalvo, 2011; Süel et al., 2006). Due to cooperativity in the amplification of siRNAs, only a sufficiently large input trigger can elicit a durable silencing response, protecting the system from aberrant noise-induced silencing. This ensures that the *unsilenced* or *ON* state is stable against small fluctuations in siRNA expression. Once a response is initiated (for example, by ingestion of dsRNA), several dynamical trajectories in the phase space of *g* and *h* are possible depending on parameter values (Figure 2D-F, see Movies S1, S2 for animations of trajectories for different regimes). Depending on the scale of the maximum amplification capacity *V*, it is possible to generate monostability (low *V*), bistability (high *V*), or an intermediate response when the system is positioned close to a saddle-node bifurcation. This latter regime, which we denote as the “critical” regime, requires fine-tuning of *V* within a very narrow parameter range, a point to which we will return presently. These three regimes make distinct predictions about the nature of the transgenerational dynamics following a trigger and, as we will argue, only a system tuned near the critical regime can explain the range of observed phenomenology summarized in Figure 1.

In the monostable regime (Figure 2D), following dsRNA exposure, the model predicts an increase in *g* followed by an increase in *h*, after which both components decay according to the respective turnover rates *γ*_1_, *γ*_2_. Generalizations of the monostable model could include possible variation between individuals due to differences in turnover rates, or asymmetric partitioning of silencing factors across and between generations as a result, for example, of small number fluctuations. However, this latter scenario would seem to be unlikely given the approximate uniformity of de-silencing across siblings, as reported by (Houri-Zeevi et al., 2020). Transgenerationally, decay is likely to be governed by a characteristic dilution rate *γ*, which corresponds to the decay of silencing factors in the absence of autocatalysis. Since the silencing factors in each animal need to be partitioned between ~250 laid eggs, one can expect that, without autocatalysis, there would be a dilution by a factor of many orders of magnitude within a memory lifetime (estimated at around a factor of 250^4^ ≈ 4 · 10^9^ after 4 generations (Houri-Zeevi et al., 2020)). This is consistent with the observation that the silencing of genes that are expressed only in the soma, and thus not amplified in the germ line, disappears after a single generation (Burton et al., 2011). This implies that 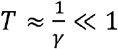 generation, while the observed average duration of silencing is found to be *T* ≈ 3 – 7 generations. This conclusion makes a monostable regime hard to reconcile with the experimental observations. Moreover, some mutants, such as the *met-2* strain, and mutants lacking piRNAs, show stable silencing for many generations, translating to hundreds of generations for the latter (Shukla et al., 2021), requiring an even lower decay rate *γ*. Finally, in the monostable setting, selection according to silencing levels would be ineffective since variation is not static, but with each generation moves directionally in phase space (Figure 2D). Based on these considerations, monostability is an unlikely candidate to explain the range of experimental phenomenology (Figure 1).

We next analyzed the regime of bistability, considered a hallmark of biological memory systems (Burrill and Silver, 2010) (Figure 2F). Here, to describe the dynamics, one may think of a thermal analogy in which the silencing state is indexed by the position of a particle in a potential well, equilibrated through interaction with its surroundings. In a monostable system, the dsRNA trigger is akin to pushing the particle to a higher position in the potential well and allowing it to relax back to equilibrium. In a bistable system, the potential well has two degenerate minima corresponding, respectively, to the ON/OFF states. In this case, a trigger that is strong enough can transition the system between the stable ON state into the adjacent metastable OFF (silenced) state. In the bistable regime, this perturbation allows the system to maintain silencing for a timescale that can far exceed that of its individual components, providing a potential explanation for long-term silencing.

However, although bistability may provide a credible explanation for some aspects of transgenerational silencing, it struggles to explain other – key – experimental observations. First, bistable mechanisms predict discrete ON/OFF states, while experiments reveal a continuum of silencing levels, such as that shown in Figure 1D (cf. Figure 2F). A second issue concerns the sensitivity of the typical memory survival time, *T*, to the underlying biochemical rates. Bistable systems implement a process of biological toggle switches, where an input stimulus transitions the system between ON/OFF states. In the *gfp* silencing experiments, while the transition from OFF to ON occurs after the presentation of the dsRNA input stimulus, the reverse transition (ON to OFF) occurs spontaneously, long after the input trigger has been removed. To address how such spontaneous transitions may occur, two explanations may be advanced: (1) there is variation in initial conditions following the removal of the trigger (see, e.g., (Houri-Zeevi et al., 2020)); or (2) temporal fluctuations can spontaneously transition the system from ON to OFF, akin to effect of thermal fluctuations on chemical reactions.

In the case of bistability with variation in initial conditions (e.g., a uniform initial distribution of *g* after triggering), the silencing state of some worms will move to the stable ON state while that of others will decay back to the stable OFF state. The former timescale could be very long, while the latter timescale is dominated by dilution and thus should decay rapidly within a single generation. Thus, overall, variation in initial conditions in the bistable regime would result in a bimodal distribution of silencing durations. This contrasts with experimental observations where intermediate silencing durations of several generations are commonly observed. Moreover, such a model would imply that the selection on a population of silenced individuals would yield a homogenous OFF population (comprising those that successfully tip over the potential barrier), contrasting with what is observed experimentally (Figure 1D). For the second possibility – bistability with noise – it is unclear how the silencing duration *T* could be set, spanning multiple generations, as the escape from the OFF state would show an extreme (exponential) sensitivity to the relative depth of the wells, the height of the barrier, and the magnitude of the noise (Kramers, 1940; Wentzell, 1998) – as emphasized by the steep dependence of *T* on *V* that we discuss below. Altogether, these considerations undermine the credibility of strong bistability as a viable model of transgenerational dynamics.

Finally, alongside monostability and bistability, there is a third scenario that is not typically considered in this context– the possibility of a time delay due to a saddle-node remnant (also known as a saddle-node *ghost*) (Strogatz, 2018) (Figure 2E), which we have referred to as the critical regime. Saddle-node ghosts are created near a saddle-node bifurcation, such as that found around the transition from a bistable to a monostable regime. In our intuitive picture of monostability and bistability, one may envisage a process (such as changing the amplitude *V*) that raises the metastable OFF (silenced) state to the level of the unstable hilltop. When the two regimes coalesce (the bifurcation), a new and nearly flat slope is formed. A particle that rolls down the slope will be delayed in this region, as if due to a “ghost” of the previous bistable fixed point. This situation occurs when system parameters are fine-tuned, for example by tuning the maximum amplification capacity *V* to a critical value *V_CRIT_*. Near the transition point, the time delay around the critical point, which will dominate the silencing memory time *T*, will have a universal behavior, in the sense that it characterizes all systems undergoing a saddle-node bifurcation near their transition point (Hathcock and Sethna, 2021). Before the transition, the delay scales in proportion to *Δ*^−1/2^, where *Δ* ≡ *V_CRIT_* – *V* is the distance from bistability (Methods). Beyond the critical point, where *Δ* < 0, the delay becomes dominated by fluctuations, with an exponential dependence on *Δ* (i.e., the delay is proportional to *e*^b|Δ|3/2^, where *b* depends on the strength of the noise – for details, see Methods). The delay can therefore become arbitrarily long, and small changes in *V* (translating to small changes in *Δ*) result in large fluctuations in average delay duration (Figure 3A).

**Figure 3.**
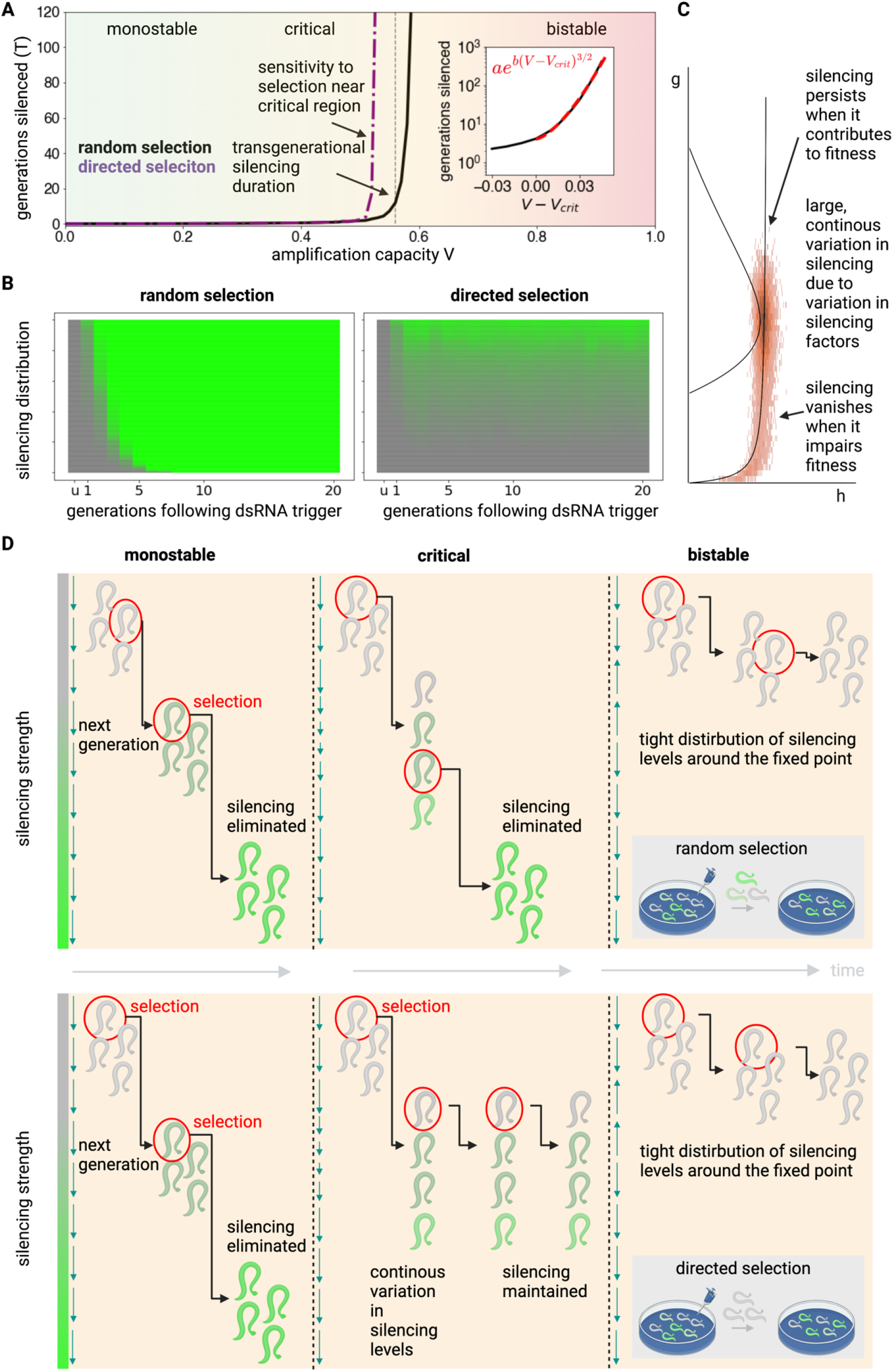
Critical configuration of the *TI* model provides long term silencing and allows selection on silencing levels. (A) Simulations of the *TI* model were performed under random selection (black) or directed selection of the most silenced individuals (purple) while varying the amplification capacity parameter *V*, exploring the phases of monostability, the critical regime and bistability. Only around the narrow critical region is there both tunability of the average silencing duration *T*, as well as high sensitivity to selection (as depicted by the large difference between *T* under the condition of directed vs. random selection). When *V* crosses the critical value *V_CRIT_* (marked by dotted vertical gray line), silencing duration begins to increase rapidly, with a stretched exponential dependence *T* ⋑ *e^a(V – V_CRIT_)3/2^* (*inset*, dashed red line, see main text and Methods for analysis). In the case of directed selection, the silencing degree s was assumed operationally to correspond to the Euclidean distance in *g, h* phasespace from the unsilenced state at (0,0), since both *g* and *h* may impact on the silencing strength. At each generation, *N* = 250 individual worms were simulated according to the *TI* model dynamics, and the subsequent generation was established by selection either from all worms (*random selection* condition) or by *directed selection* of worms with *s* > *s_select_*, corresponding to strong silencing (here, we took *s_select_* = 3 and set the generation time as 72 hours). (B) Silencing distributions over time in the case of either random selection (left panel) or directed selection (right panel) (Methods). (C) The large, static, variation around the critical point allows for selection according to silencing levels. Simulation details are the same as in Figure 2E, where the variation represents the distribution of silencing factors after a single generation starting from above the critical point. All simulations used the same parameters as in *Table 1*, where *V* is set so the system lies in the critical regime. The critical regime yields a stationary, continuous, distribution of silencing states, as observed by (Vastenhouw et al., 2006) (cf. Figure 1D). (D) Schematic illustration of the differential effects of random (upper panel) vs. directed (lower panel) selection on silencing dynamics in the three major regimes of the system: monostability, the critical regime and bistability. In the critical regime, due to large static variation around the bifurcation point, there is high differential sensitivity to selection.

In contrast to the monostable and bistable regimes, the critical regime can provide a credible explanation of the full range of experimental phenomenology described in Figure 1. The silencing memory duration, *T*, depends on *Δ* and, as such, can be effectively decoupled from the turnover rates of the underlying molecular components. Small fluctuations in *Δ* translate to large differences in silencing times between individuals, characterized by a survival distribution with a nearly flat tail (Methods). When silenced, the silencing state will become “delayed”, and therefore concentrated, near the critical region. Here, the system becomes subject to the phenomenon of *critical slowing down* (Scheffer et al., 2009), where the rate of change of *g* and *h* becomes very small, and the variance (e.g., of the silencing levels) between individual trajectories (worms) increases. Critical slowing down therefore results in large static variation between individuals of the same generation, and the emergence of a continuum of silencing states (Figure 2E, lower panel), contrasting the existence of binary OFF/ON states expected for a mechanism based on bistability. Finally, the model predicts little inter-individual variation at the final de-silencing stage, as this region corresponds to the phase of monostable decay after the ghost region has been traversed, reducing variation between de-silenced sisters.

The phenomenon of critical slowing near the bifurcation endows the system with high sensitivity to selection according to silencing levels (Figure 3, see Movie S3 for animation of a population of stochastic trajectories, and Movie S4 for an animation of deterministic and random selection). When individuals are selected at random at each generation, progression of the population in phase-space proceeds with a typical delay around the critical region, with a timescale of several generations (Figure 3A, B – left panel, C, D – upper panel). By contrast, directed selection of individual worms in which genes are most strongly silenced allows silencing to be maintained for much longer (and potentially indefinite) periods of time (Figure 3A, B – right panel, C, D – lower panel): This phenomenon occurs due to the combination of delayed or “static” progression and large variance of the population in the critical region, which allows for selection according to silencing levels. Recurrent selection of silenced worms on this background leads to the emergence of a persistent distribution of silencing levels in their progenies away from the “de-silenced” state, as observed experimentally (compare Figure 3B to Figure 1D). This contrasts with the monostable regime, where selection is ineffective due to the monotonic progression in phase-space, and the bistable regime, where variation between individuals is suppressed near the stable point. Taken together, the ghost (critical) mechanism provides a plausible timer mechanism for transgenerational inheritance of silencing in *C. elegans.*

### Robust tuning of transgenerational silencing memory persistence is a direct result of competition between genes over silencing resources

Despite its success at explaining the observed experimental phenomenology, a major limitation of the ghost mechanism is its apparent reliance on the fine-tuning of the gap *Δ*, the distance of the model parameters from the point of saddle-node bifurcation, to set the silencing duration *T.* Small deviations in *Δ* lead either to a very short silencing duration *T* (cf. a monostable regime) or very large *T* (a bistable regime), as illustrated in Figure 3A. The mechanism is therefore not robust – noise and variation in circuit parameters can easily change the timescale *T* by orders of magnitude.

At first sight, it is unclear how a biological system can tune and maintain *Δ* in a robust manner. One possibility is inspired by the concept of self-organized criticality in statistical physics where, under certain conditions, a ghost regime *(Δ* ≈ 0)can emerge as an attractor state (Bak et al., 1988; Vidiella et al., 2021), rather than due to fine-tuning of a set of fixed parameters. A classic conceptual example was developed by Bak et al., where a sandpile is formed by slowly adding grains of dry sand (Bak et al., 1988). Over time, such a system settles around a critical state, where the addition of new sand grains must, on average, be balanced by the removal of others through avalanches (Bak et al., 1988; Frette et al., 1996). Indeed, the concept of such self-organized criticality has been applied to biological systems in multiple contexts, including genetic networks, brain activity and animal flocks (Mora and Bialek, 2011; Moreau et al., 2003; Muñoz, 2018), including in particular a classic study on self-tuning to a Hopf bifurcation in the context of hearing (Camalet et al., 2000; Eguíluz et al., 2000). More recently, Stanoev et al. have proposed that receptor activity could become self-organized at a saddle-node ghost through fluctuation sensing (Stanoev et al., 2020). However, by itself, self-tuning to criticality *(Δ ≈* 0) does not present a robust mechanism to tune the timescale *T*, which requires precise tuning of *Δ* in the vicinity of the critical point.

How, then, is such an apparently fine-tuned regime achieved? In the following, we will show that selforganization of the ghost regime, in fact, emerges as a consequence of a key feature of the RNA-mediated transcriptional silencing machinery in *C. elegans*, the competition for limited amplification resources (as noted in Figure 1F). Moreover, within the framework of a minimal competition model, we will see that the average inheritance timescale, *T*, can be set robustly, even in the presence of noise and variation between genes.

In order to account for competition between siRNAs, we developed a straightforward generalization of the *TI* model, which we termed the *Toggle-Inhibitor Competition (TIC*) model. While competition has been considered previously in mathematical models of transgenerational inheritance (Houri-Ze’evi et al., 2016), these theories, which only account for competition between two silencing pathways, cannot explain the observed tuning. Here, we show that a “many-particle” theory, that accounts for competition between many silenced genes, provides a robust mechanism of fine-tuning. Let us denote by indices *i* = 1,…, *N* the ensemble of endogenous and exogenous genes that can become transgenerationally silenced. Silencing events occur stochastically, following from either chance encounters with exogenous challenges such as viruses, or endogenous stochastic silencing events (Beltran et al., 2020). Within the framework of the TIC model, we propose a model of silencing dynamics similar to that of the *TI* model but with a global “cost” term *C*(*g*_1_, *h*_1_, *g*_2_, *h*_2_,..), which inhibits the amplification rate of siRNAs by modulating *V* or *k*_1_ (see Movie S5 for the effects of modulation *V* or *k*_1_ on the nullclines). Here, for simplicity, we begin by supposing that the model parameters, *k*_1_, *k*_2_, etc. do not vary between genes. To demonstrate the role of resource competition in the TIC model, we will consider a simple scenario in which the cost function modulates the amplitude 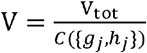 with *C* = ∑*_j_g_j_* noting that other possibilities, including modulation of *k_1_* or other choices of *C* lead to similar dynamics (Methods). In this case, the TIC model translates to the kinetics

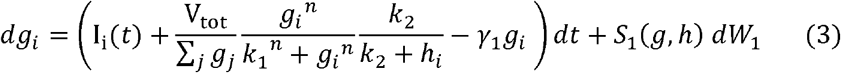

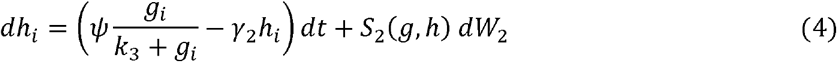

where, in this case, *I_i_*(*t*) is a stochastic random variable. For example, I_i_(*t*) may acquire positive values at a constant rate, so that the number of silencing events per generation is characterized by a Poisson distribution. Note that we have also replaced the amplitude *V* by the parameter *V*_tot_, as this now denotes the amplification capacity that is shared between genes.

For simple circuits consisting of linear production and removal, competition for synthesis resource would reduce the overall activity level, leading to variation in the steady-state or amplitude without affecting response duration. However, for the *TIC* model, a strikingly different dynamical pattern emerges (Figure 4). Consider a situation in which the system begins in a state with no silenced genes, so that the initial cost function *C* is small and the amplification capacity large, placing the system in a phase of bistability. Under these conditions, silenced genes will begin to accumulate through stochastic silencing events, leading in turn to a gradual increase in the cost function *C*. Stochastic simulations of the model dynamics are depicted in Figure 4A,B, as well as Movie S6 in animation. If this increase occurs rapidly, the cost function may increase to a level that places the system in a monostable regime, where genes now become rapidly de-silenced, leading in turn to a decrease in *C.* Based on these dynamics, this process will continue until the system eventually stabilizes close to the region of bifurcation between bistability and monostability, where the ghost regime is defined (Figure 4B). Of course, in practice, convergence to this phase from the bistable regime may not be characterized by such a damped oscillatory dynamic, but the approach may be monotonic. Yet, within this framework, the stability of the critical ghost regime in the long-term is assured with no requirement for fine-tuning of biochemical parameters.

**Figure 4.**
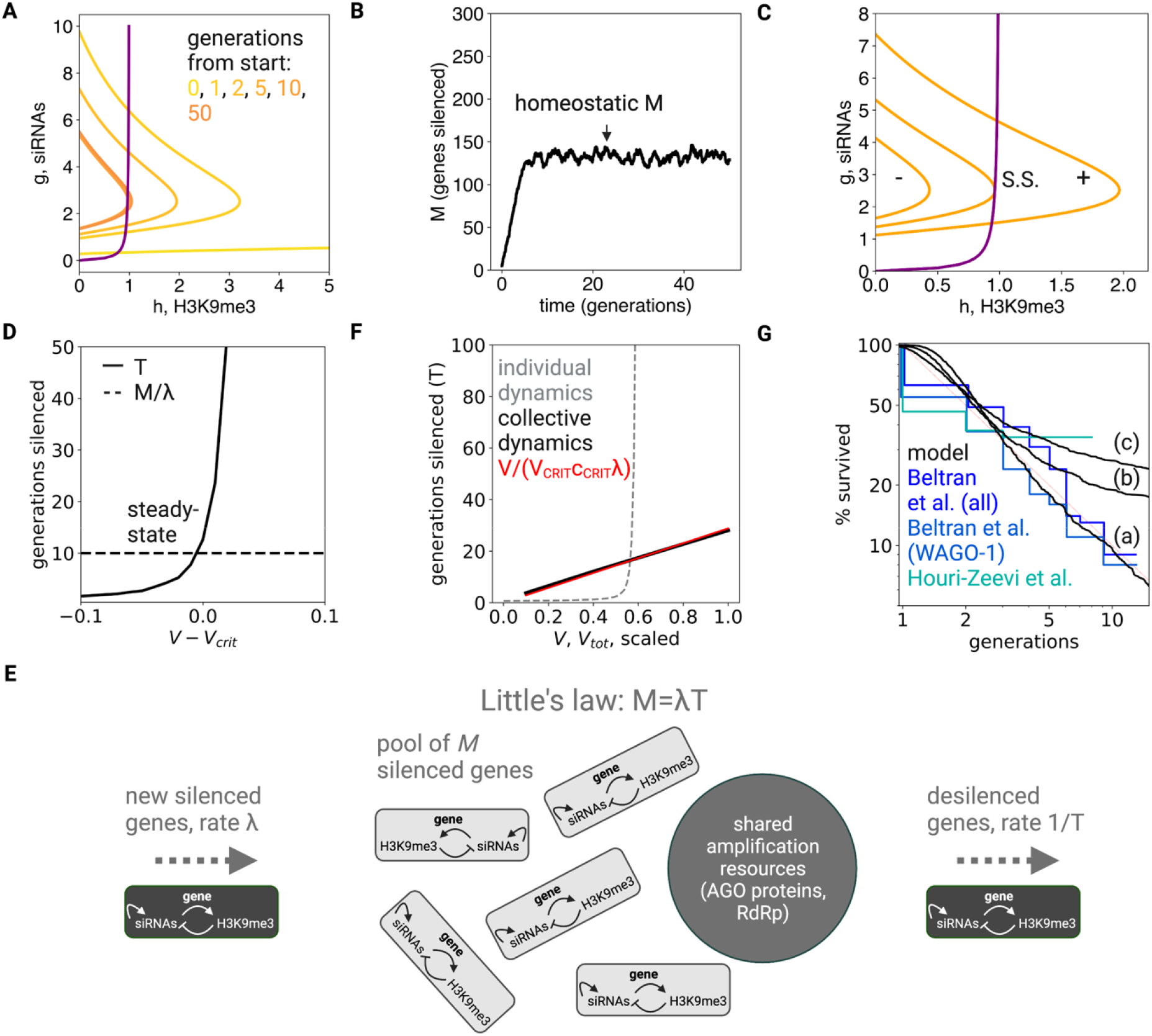
Robust self-organization of long-term gene silencing driven by competition for silencing machinery. In the *Toggle-Inhibitor-Competition (TIC)* model, stochastic events in a pool of candidate genes drives silencing transitions. We denote by the average number of silenced genes at steady-state, and by λ the average rate at which new silencing transitions occur stochastically. (A) Starting with an initial condition in which all genes are unsilenced, stochastic simulation of the *TIC* model shows nullclines (purple) and (orange, different lines depict generation number) settling near the region of saddle-node bifurcation over time. Note the rapid convergence from bistability towards the critical regime between bistability and monostability. (B) Over time, the system settles into a stable homeostatic phase in which, at any given time, an average of *M* genes are silenced. (C) The configuration of nullclines *ḣ* = 0 (purple) and *ġ* = 0 (orange, different lines for monostable, ghost and bistable configurations) set by the bifurcation ratio of the amplitude and cost function *V*_tot_/*C*, determines the average silencing duration time, *T.* When the system is in the bistable regime (right-most orange nullcline), silenced genes accumulate (+); while in the monostable regime (left-most orange nullcline), they are removed (-). There is thus feedback from the size of the silenced gene pool *M* back to the bifurcation parameter through the cost term driving the system to steady-state (S.S.). (D,E) Schematic illustration of the mechanism underlying the tuning of *T.* According to Little’s law (depicted graphically in panel E), at steady-state, *M* = *λT*. Note that, while the silencing duration *T* (panel D) changes rapidly around the bifurcation point (results from stochastic simulation as in Figure 3A), *M* is much more narrowly distributed (panel B), fixed by the cost function near the bifurcation. This narrow range of *M* effectively sets the average duration time *T* = *M*/*λ*. (F) The silencing duration *T* is proportional to *V*_tot_ (black line, results of stochastic simulation while varying *V*_tot_) and can be recovered by a simple analytical formula (red line, see main text and Methods). This behavior contrasts with the high (exponential) sensitivity of silencing duration on the amplification parameter *V* in the *TI* model (gray line, stochastic simulation of *TI* model while varying *V*). (G) Variation around the critical point leads to a heavy-tailed distribution of silencing durations, due to the high sensitivity of silencing durations around that point. Silencing duration distributions were generated from simulations of the *TI* model *V* drawn from a normal distribution with mean 〈*V*〉 = 0.55 and (a) 4%, (b) 10%, (c) 15% variation around the mean. Simulation parameters are provided in *Table 2*, and simulation details and code are provided in Methods.

To understand why the system must settle near the saddle-node bifurcation, it is instructive to consider the dynamics of the system from the perspective of stochastic queuing theory (Figure 4C-E). Here, the siRNA silencing machinery can be considered as a “server” and silenced genes as “clients” that arrive at random and are handled by the server before becoming de-silenced. If *M* denotes the average number of genes that are silenced when the system is in steady-state, from the intuition above, it follows that the system can be bistable at steady-state only when *M* = *N*, the total number of genes that can be silenced. This situation will prevail only when the amplitude V_tot_ is sufficiently large. However, at smaller values of V_tot_, *M* « *N*, and the system must transition to the critical phase of monostability long before all genes become silenced. In this case, if we denote by *λ* the average arrival rate of new silencing events (a property of the stochastic input rate *I_i_*(*t*)), at steady-state the average silencing duration would be given by *T* = *M*/*λ*, a general property that follows from Little’s law for the steady-state dependency of a queuing system (Little, 1961). As the system transitions stochastically between bistability to monostability, varies around a narrow range (due to small changes in the cost parameter), while silencing durations of individual genes cover a wide range of possible values. The system thus fixes the average gene silencing duration according to the relation in the critical region. We note that, even in the presence of noise, which has the effect of pushing the system marginally towards bistability, the system will settle near the bifurcation due to the very steep (exponential) dependence of on the amplitude (as depicted in Figure 4D, see Methods).

### Quantitative properties and predictions of self-tuned criticality by competition

The *TIC* model provides a robust mechanism for tuning the average silencing duration *T* independently from the individual turnover rates of the underlying molecular components. The capacity *M*, the average number of silenced genes at any given time, can be derived directly from the model – it is fixed by the average cost per silenced gene near the bifurcation point c_CRIT_, and is proportional to the maximal amplification capacity, *V*_tot_/*c*_CRIT_ (Methods; note that, here, *c*_CRIT_ is simply *g*_CRIT_, the value of *g* at the critical point). This holds even when considering noise or statistical variation in circuit parameters between genes (Methods). We can therefore think of the *TIC* model as defining a mechanism that translates the amplification capacity *V*_tot_ into an average silencing memory time *T* (Figure 4E,F). Indeed, this mechanism can explain the association between hypersensitive RNAi responses and long-term silencing in *met-2* mutants (Lev et al., 2019).

Crucially, the *TIC* model predicts a gradual (linear) dependence of *T* on *V*_tot_ and other circuit parameters, which contrasts with the steep dependence of *T* on *V* for individual memory dynamics near the critical point (Figure 4E,F). This linear dependence can be tested by experimentally manipulating *V*_tot_. The linear dependence of the average memory time on *V*_tot_ allows for robust tuning of silencing duration since changes in *V*_tot_ only lead to proportional changes in *T*. This allows for regulatory programs and evolutionary pressures to robustly tune and adjust a “forgetting time” on the order of many generations.

The change in *T* following a change in circuit parameters is due to the collective dynamics of the silencing memories. For example, a change in *V*_tot_ changes the size of the steady-state silenced gene pool *M* (*M* → *M*′). The dynamics of the change may be asymmetric in their dependences: A downwards change in *M* shifts transiently the system into the monostable regime, leading to rapid de-silencing. On the other hand, depending on the scale of Λ, an upwards change in *M* increases the number of silenced genes over a time scale on the order of *T* = *M*′/*λ* (Methods). Similar dynamics also hold for an individual silenced gene, which upon a downwards change in *M* may become de-silenced rapidly; but upon an upwards change, will become re-silenced within a typical timescale of *N*/*λ* > *T*, under the minimal assumption that all genes are silenced at the same rate. Such slow recovery from a downwards to upwards change is evident in an experimental study by Klosin et al. (Klosin et al., 2017). Animals expressing a stochastically-silenced multi-copy reporter transgene were raised in high temperature conditions (25 °C), which causes the de-silencing of endogenous genes (viz. a step change in some physiological parameters). Re-silencing can be established by transferring the worms back to 20 °C (reversing the previous step change). Klosin et al. demonstrated that a mere 48-hour incubation in 25 °C caused a de-silencing that persisted for 7 generations, while a 5-generation incubation in 25 °C caused de-silencing that persisted for 14 generations. In the model, the slow dynamics of recovery reflects the stochastic nature of the accumulation process.

The linear dependence of *T* on *V*_tot_ is a property of the system when the silencing pool is near steadystate, while perturbations in *V*_tot_ may result in transient monostable or bistable behavior. The silencing time *T* is thus predicted to show transient history dependence with a multi-generational timescale. Bistability may become permanent if *V*_tot_ (or other model parameters) change so that *M* ≈ *N*; that is, the steady-state silenced gene pool contains all genes that can become silenced.

A final prediction concerns the distribution of silencing durations. The time *T* is an average statistic, and there can be large variations due to stochastic fluctuations in circuit parameters, as well as variation between genes, which are typical of biological systems (see Movie S7 for simulation of the TIC model with variation in parameters between genes). Since the system is positioned close to a saddle-node bifurcation, the silencing dynamics of an individual gene may be positioned at some distance *δ* from the bifurcation, and we can consider *δ* as a random variable. When *δ* < 0, silencing time scales are short, generating a steep “head” of silencing durations with a distribution that decays as 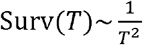, while, for *δ* > 0, silencing durations are much longer (as individual genes are in their bistable regime), generating a heavy tail that may approach a nearly-uniform survival function (Figure 4G and Methods). Together, these dependences provide an explanation for the heavy-tails shown in the silencing time distribution (Figure 4G). It is important to note that, due to universal scaling near the critical point, these predictions are independent of model details and are only due to the phenomenon of self-organization. They are thus predicted to characterize general systems that tune near to a saddle-node bifurcation through collective dynamics.

Together, the *TIC* model suggests that long-term memory in the *C. elegans* silencing system arises from the collective dynamics of many individual memory (gene silencing) events that compete over shared synthesis resources. This process of mutual competition causes the dynamics of the entire system to self-organize near a critical point (a saddle-node ghost), where individual silencing events become delayed. We thus term the mechanism *self-tuned criticality by competition*. The delay near the critical point may be much longer than the turnover times of the underlying molecular machinery, and so determines the average memory duration. Additionally, self-organization near the critical point yields a static and continuous distribution of silencing strengths that can be selected upon.

## Discussion

Epigenetic gene-silencing in *C. elegans* identifies a general design principle for the implementation of long-term memory by biological circuits. We showed that the wide range of experimental phenomenology may be captured by a response system positioned close to a saddle-node bifurcation. By combining a mathematical modelling-based approach with transcriptional and chromatin measurements, we found that the dynamics of gene silencing are consistent with an excitable circuit model, where silencing is driven by a bistable switch, followed by inhibition. Competition between silencing memories for limited resources self-tunes the response system near to a saddle-node bifurcation. These core interactions, encapsulated by the *Toggle-Inhibitor-Competition* model, provide a robust timer mechanism for transgenerational silencing dynamics.

The *TIC* model contrasts with classical models of biological memory. From the perspective of biological circuits, long-term memory is often attributed to bistable switches, which can transition between stable ON/OFF states (Alon, 2019; Burrill and Silver, 2010; Ferrell, 2021). Bistable switches arise in systems with nonlinear positive feedback (Ferrell and Xiong, 2001; Ninfa and Mayo, 2004). An activating stimulus can shift the system from the OFF to the ON state, and the system will remain in the ON state even after the stimulus has been removed. Bistable switching is thought to stand at the core of a myriad of biological processes, including the control of cell cycle transitions (Doncic et al., 2015; Pomerening et al., 2003), cellular differentiation (Wang et al., 2009), vernalization (Angel et al., 2011), and immune cell homeostasis (Hart et al., 2014). By contrast, in the *TIC* model, although activation dynamics are governed by bistability, the system becomes self-tuned to the vicinity of a critical regime, and the delay in this regime determines memory lifetime.

While bistable switches allow biological systems to maintain memory, they have several drawbacks that hinder their applicability in memory systems such as the transgenerational inheritance system. The first issue concerns the maintenance of a robust “forgetting time” (or survival time) *T* that is neither too short nor too long. Forgetting requires spontaneous transitions between states in the absence of an explicit trigger, leading to a typical exponential dependence on both the magnitude of noise and circuit parameters (as exemplified by the escape rate from a potential well for gradient systems, but which can be extended to non-gradient systems via large deviations theory) (Kramers, 1940; Wentzell, 1998). Such exponential dependence is, however, not robust as it corresponds to a high sensitivity of *T* to specific cellular conditions.

The second issue concerns the ability to retain certain relevant memories over other memories, according to inputs to the system. This is exemplified in the transgenerational silencing system by the sensitivity to selection on silencing levels – the critical regime endows the system with a continuum of nearly static silencing levels, which allows for efficient selection. This contrasts with the monostable regime, where trajectories advance in phase-space with each generation, and the bistable regime, where fluctuations decay rapidly towards steady-state. In the critical regime, selection of worms with stronger silencing provides a perturbation that prevents average silencing from decaying away from the critical point. More generally, one can consider other perturbations that move the memory state in phase space, including continuous input signals. As an example, in the epigenetic memory system, repeated priming resulted in extension of silencing retention (Houri-Ze’evi et al., 2016). Self-organization near the critical point provides a mechanism for integrating these perturbations in order decide on memory retention (Stanoev et al., 2020).

The TIC mechanism decouples the average memory duration *T* from the underlying molecular dynamics. According to the model, *T* is given by the ratio of the size of the memory pool *M* (fixed by the location of the critical point in phase-space) to the arrival rate of gene silencing events *λ*. Indeed, it is also possible to decouple *T* from *λ* by adding a feed-forward activation of synthesis resources *V* by memory arrivals, as this would normalize away *λ* from *T* (Methods).

In this study, we proposed a simple realization of the *TIC* model based on autocatalysis of target-specific siRNAs and negative feedback from silencing chromatin marks. While such a model is plausible based on the known biological interactions and dynamical measurements, there are likely to be other important regulatory interactions that affect transgenerational inheritance dynamics. However, due to the generality of the model assumptions, as well as scaling behavior near the critical point, the important quantitative aspects of the model are preserved even in more complex and higher-dimensional settings. For example, one may show that the nature of the dynamics is unchanged when including into the dynamical equations pUG RNA-templates from which siRNAs are synthesized ((Shukla et al., 2020), Methods). Moreover, the observation that competition over synthesis resources leads to self-organization near a critical point allows us to make quantitative predictions on the properties of the system (such as the distribution of memory durations) despite having only limited knowledge of the underlying regulatory network.

Given the simplicity of the model, it makes sense to question whether the concept of self-tuned criticality by competition can be extended to other memory systems, including cell-based memory systems such as the mammalian immune system, as well as cellular interaction networks (Figure 5). We, therefore, summarize the few necessary ingredients that such a memory system should possess, and the signature properties that may indicate that such a mechanism is at play. Specifically, for the mechanism to operate, the system needs to have memory response dynamics governed by an excitable circuit, with cooperative activation and negative feedback that are memory-specific. In the transgenerational silencing system, these memory-specific interactions may arise by siRNAs that silence the same target sequence. In cellular systems, these may correspond to intracellular excitable circuits; indeed, such circuits have been implicated in the process of cellular differentiation (Chang et al., 2008; Kalmar et al., 2009). In addition to the memory-specific excitable circuit, the model also requires *global inhibition*, which can act as a control parameter for the transition from monostability to bistability in individual memories. Global inhibition can occur in different flavors. However, the simplest form is inhibition of the amplification rate through competition over synthesis resources. In the case of the *C. elegans* inheritance system, there is competition of AGO proteins and RdRP complexes. In the immune system, T-cells compete over global factors such *IL-7* that control overall T-cell levels (Bradley et al., 2005), while other factors such as *FGF* play an important role in mediating quorum feedback in cellular dynamics (Kitadate et al., 2019; Saiz et al., 2020). Such mechanisms are widely regarded in the biological literature as *homeostatic*, as they maintain the overall levels of memory units within a fairly narrow range. Finally, it is required that the average memory timescale *T* be longer than the typical timescale of the turnover rate of the excitable memory circuit. The combination of fast excitable memory-specific dynamics with global inhibition keeps the system near bifurcation by pushing it away from bistability and allows for robust tuning of memory duration *T.*

**Figure 5.**
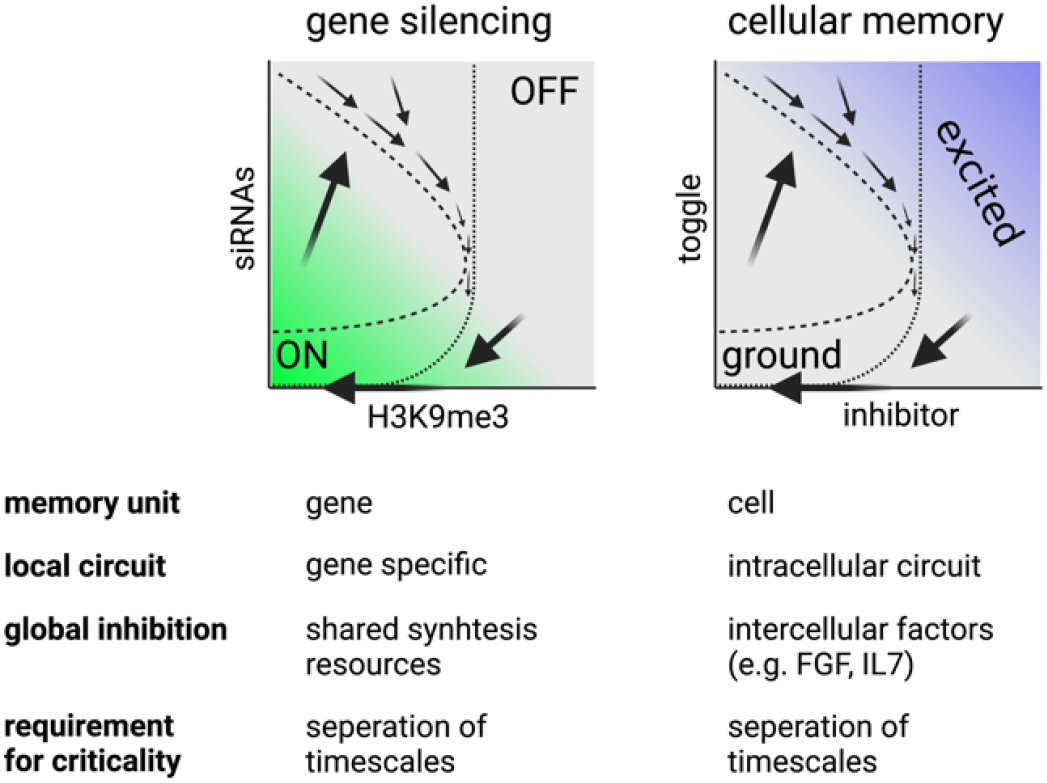
Generalization of self-tuned criticality by competition to cell-based memory. The *TIC* model can be generalized to cell-based memory circuits, where the “local” memory unit is a cell (compared with a gene for the silencing mechanism in *C. elegans).* The model requires an intracellular excitable circuit separating “ground” and “excited” states, similar to the excitable high/low Nanog circuit described by Kalmar et al. in the context of embryonic stem cells (Kalmar et al., 2009). In the immune system, the excited state may correspond to a memory identity associated with a particular antigen. It also requires global feedback that can transition the system between monostable and bistable states, which may be implemented by known quorum sensing mechanisms. Finally, self-organization near the critical state is achieved when there is a separation of timescales between the steady-state memory duration and the turnover rates of the underlying molecular circuit.

The model makes several key predictions that can serve as a signature when considering its potential applicability in other memory systems. First, in the framework of TIC dynamics, memory duration depends on collective memory dynamics. In an “open loop” setting, where collective feedback is not active, memory duration is predicted to show very high sensitivity to circuit parameters around the critical point while, in the “closed loop” setting, self-organization around the critical point leads to only a mild dependence of on circuit parameters. Second, according to the model, memory duration may be adjusted by changing the memory pool size by adjusting synthesis capacity, or by modulating the arrival rate. Moreover, the average memory duration can be inferred by examining the time until the re-establishment of the memory repertoire from stochastic arrival events. Finally, the model predicts a distinct distribution of memory survival times that results from its convergence at steady-state towards the vicinity of a saddle-node bifurcation (Methods), with a relatively steep head corresponding to events taking place closer to the monostable regime, and a flat tail (that asymptotes towards a nearly flat dependence on) corresponding to events taking place closer to the bistable regime.

While the transgenerational silencing of endogenous and exogenous genes in *C. elegans* is well established, the evolutionary benefits of the mechanism remain a topic of intensive research, with a focus on adaptation to fluctuating environments and for pathogen memory (Frolows and Ashe, 2021). A particularly intriguing aspect is the widespread stochastic silencing of genes (Beltran et al., 2020). Stochastic, transient silencing provides a mechanism for stochastic phenotypic switching, which can have several functional benefits such as bet-hedging against environmental uncertainty (Kussell and Leibler, 2005; Silva et al., 2021; Veening et al., 2008; Xue and Leibler, 2017). Our model provides a generic mechanism for the implementation of phenotypic switching over intermediate timescales. Moreover, rather than binary switching between discrete states, our model suggests that silencing at the population level can be maintained at a heritable continuum. Thus, if a rare stochastic silencing event provides an unexpected benefit (such that stronger silencing is associated with higher fitness), it can be maintained for a longer duration, while detrimental silencing events are discarded. This may allow worms to adjust and adapt their pattern of gene expression in response to changing environmental conditions.

In conclusion, competition in the TIC model provides a robust mechanism for tuning long-term memory persistence and presents features beneficial for the retainment of relevant memories.

## Data and code availability

The codes for stochastic simulations presented in this study are available from the corresponding authors upon request and will be made openly accessible on public depositories at the time of publication.

## Acknowledgements

We are grateful to Peter Sarkies (University of Oxford), Yael Korem (Yale University), and members of the Simons lab for useful discussions. O.K. acknowledges support from the James S. McDonnell Foundation 21st Century Science Initiative Understanding Dynamic and Multi-scale Systems - Postdoctoral Fellowship Award. This work was supported by grants to B.D.S. (Wellcome Trust 219478/Z/19/Z and the Royal Society EP Abraham Research Professorship, RP/R1/180165). The research team also acknowledges core support from the Wellcome Trust (092096) and CRUK (C6946/A14492).

## Author contributions

O.K., E.A.M. and B.D.S. conceived the project. O.K. formulated, analyzed and interpreted the TI model and its generalization with continual input and advice from E.A.M. and B.D.S. O.K. and B.D.S. wrote the manuscript with further input from E.A.M.

## Declaration of interests

The authors declare no competing interests.

## Open Access Requirements

This research was funded in whole, or in part, by the Wellcome Trust (098357/Z/12/Z and 219478/Z/19/Z). For the purpose of Open Access, the author has applied a CC BY public copyright license to any Author Accepted Manuscript version arising from this submission.

## Notes

### Competing Interest Statement

The authors have declared no competing interest.

