## Supplementary Information for "Epigenetic inheritance of gene-silencing is maintained by a self-tuning mechanism based on resource competition"

### Methods

In the following sections, we provide further details on the Toggle-Inhibitor (TI) model of gene silencing and its generalisation, the Toggle-Inhibitor-Competition (TIC) model. Specifically, we use analytical approaches to determine the properties of these models including their phase behavior, stability, dynamics and the effects of noise, as well as presenting details of the stochastic simulations used to model the different conditions considered in experiment.

#### 1 Toggle-Inhibitor (TI) model

The TI model is based on excitable dynamics with an effector $g$ (the concentration of gene-specific 22G siRNAs) and an inhibitor $h$ (the concentration of silencing marks H3K9me3 on the target gene). In the absence of noise, its dynamics are given by the coupled set of rate equations, Eqs. (1,2) of the main text:

$$\dot{g} = I(t) + V\frac{g^n}{k_1^n + g^n}\frac{k_2}{k_2 + h} - \gamma_1 g \tag{1}$$

$$\dot{h} = \psi\frac{g}{k_3 + g} - \gamma_2 h \tag{2}$$

where $I(t)$ denotes the source term or trigger. Since both $g$ and $h$ contribute to silencing, we can consider the degree of silencing $s$ to be an increasing function of $g$ and $h$, such as the Euclidean distance from the origin: $s = \sqrt{g^2 + h^2}$. The effector is catalyzed through cooperative autocatalysis with a Hill-coefficient $n > 1$, maximal synthesis rate $V$, and half-way saturation $k_1$. Synthesis is inhibited by $h$, with a half-way inhibition $k_2$. There is also a production term (e.g., due to dsRNA triggers) set by $I(t)$. The inhibitor is synthesized by $g$ with half-way saturation $k_3$ and maximal amplification rate $\psi$. The effector $g$ decays with rate $\gamma_1$ and the inhibitor $h$ decays with rate $\gamma_2$. For simplicity, we consider all the concentration variables $k_1, k_2, k_3$ as defined in arbitrary units of concentration (denoted AU).

The TI model captures the dynamics of silencing factors on a multi-generational timescale, and therefore incorporates the effect of dilution due to growth and reproduction. We denote by $\tau$ the time scale of a single generation, which is on the order of days (see Table 3). Time scales can thus be divided into generations, starting from $t = 0$ (first generation, second generation, etc.). For the rate parameters, we note that the dynamics on the transgenerational timescale are dominated by dilution, and a reasonable estimate for the decay rates $\gamma_1$, $\gamma_2$ would be on the order of hours (see main text).

#### TI model nullclines

To understand the dynamics of Eqs. (1,2), we first consider their nullclines, where $\dot{g} = 0$ or $\dot{h} = 0$. For simplicity, we will start by setting $I(t) = 0$, noting that a non-zero production rate changes the nullclines (a situation that we will analyze later). The nullclines for Eq. (1) are given by:

$$g = 0 \tag{3}$$

$$h = k_2 \left( \frac{V}{\gamma_1} \frac{g^{n-1}}{(g^n + k_1^n)} - 1 \right) \tag{4}$$

while, for Eq. (2):

$$h = \frac{\psi}{\gamma_2} \frac{g}{k_3 + g} \tag{5}$$

The nullclines are illustrated graphically in Figure S1 for sample parameters. The nullcline of $\dot{h}$ is a Michaelis-Menten curve, which saturates at a maximum $h = \psi/\gamma_2$, while the non-trivial nullcline of $\dot{g}$ is a unimodal curve with an extremum at position $p_{\max} = (g_{\max}, h_{\max})$ where, solving $\partial h/\partial g = 0$ on the nullcline, one finds that

$$g_{\max} = (n-1)^{1/n} k_1 \approx k_1, \qquad h_{\max} \approx \left( \frac{V}{2\gamma_1 k_1} - 1 \right) k_2 \tag{6}$$

Thus, for $V/\gamma_1 < 2k_1$, the non-trivial nullcline takes values only at negative $h$, and only the trivial nullcline is relevant. Note also that, for $g \gg k_1$, the nullcline decays approximately linearly as $h \approx k_2(V/\gamma_1 g - 1)$.

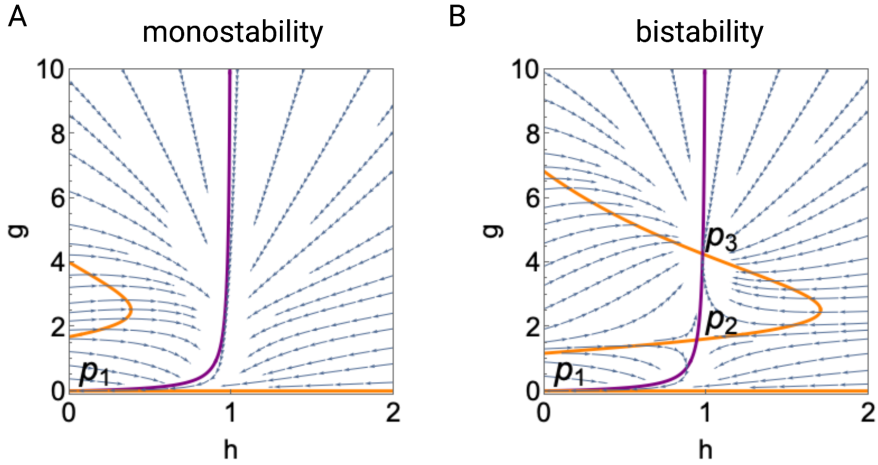


**Figure S1:** Stream plots of Eqs. (1,2) and nullclines for $\dot{g} = 0$ (orange) and $\dot{h} = 0$ (purple), with parameters as in Table 1, and (A) $V = 0.45$ AU/hr or (B) $V = 0.7$ AU/hr. For (A), the nullclines intersect at $p_1 = (0, 0)$ placing the system in a monostable regime. For (B), the nullclines intersect at three points, $p_1$, $p_2$ and $p_3$, placing the system in the regime of bistability.

#### Fixed points and stability

The fixed points of the system are given by the intersections of the nullclines of Eqs. (1,2). For the trivial nullcline $g = 0$ there is one intersection at $p_1 = (0, 0)$. To assess the stability of this fixed point, we can use linear stability analysis, forming the Jacobian,

$$\mathbf{J} = \begin{bmatrix} \partial_g \dot{g} & \partial_h \dot{g} \\ \partial_g \dot{h} & \partial_h \dot{h} \end{bmatrix} = \begin{bmatrix} V \frac{n g^{n-1} k_1^n}{(k_1^n + g^n)^2} \frac{k_2}{k_2 + h} - \gamma_1 & -V \frac{g^n}{k_1^n + g^n} \frac{k_2}{(k_2 + h)^2} \\ \psi \frac{k_3}{(k_3 + g)^2} & -\gamma_2 \end{bmatrix} \tag{7}$$

At the fixed point $p_1$, the Jacobian

$$\mathbf{J}(p_1) = \begin{bmatrix} -\gamma_1 & 0 \\ \frac{\psi}{k_3} & -\gamma_2 \end{bmatrix} \tag{8}$$

has two negative eigenvalues $l_1 = -\gamma_1$, $l_2 = -\gamma_2$, and is therefore stable, with decay of the state towards $p_1$ dominated by dilution.

The existence of other fixed points depends on the location of $p_{\max}$ and whether the non-trivial nullcline of $\dot{g}$ intersects the nullcline of $\dot{h}$. There may be zero, one, or two intersection points. Here, we will analyse the case where the synthesis rate is large, so there are two intersection points (see Figure S1). Taking $g \gg k_1$, the first intersection point, which we denote as $p_2 = (g_{p_2}, h_{p_2})$, occurs at $g_{p_2} \approx \frac{\sqrt{\gamma_2 k_2 V}\sqrt{4\gamma_1 k_3 \psi + \gamma_2 k_2 V} + \gamma_2 k_2 V}{2\gamma_1 \psi} \approx \frac{\gamma_2 k_2}{2\gamma_1 \psi} V$ with $h_{p_2}$ obtained from Eq. (5). Then, noting that

$$\begin{aligned} V\frac{n g^{n-1} k_1^n}{(g^n + k_1^n)^2}\frac{k_2}{k_2 + h_{p_2}} &\approx \frac{k_2 n V g^{-n-1} k_1^n}{h_{p_2} + k_2} \to 0 \\ -\frac{k_2 V g^n}{(h_{p_2} + k_2)^2 (g^n + k_1^n)} &\approx -V\frac{g^n}{k_1^n + g^n}\frac{k_2}{(k_2 + h_{p_2})^2} = C_1 V \\ \frac{k_3 \psi}{(g + k_3)^2} &\approx \frac{4\gamma_1^2 k_3 \psi^3}{\gamma_2^2 k_2^2 V^2} = C_2 V^{-2} \end{aligned}$$

the Jacobian at $p_2$ is given approximately by

$$\mathbf{J}(p_2) \approx \begin{bmatrix} -\gamma_1 & C_1 V \\ C_2 V^{-2} & -\gamma_2 \end{bmatrix} \tag{9}$$

In this case, noting that $V \gg 1$, the Jacobian has eigenvalues $l_1 \approx -\gamma_1$ and $l_2 \approx -\gamma_2$. Since both eigenvalues are negative, the fixed point $p_2$ is also stable, with the system decay rate dominated by the decay rates $\gamma_1, \gamma_2$. Finally, the other intersection point $p_3$ occurs at a concentration $g_{p_3}$ below $k_1$, where the term $V\frac{n g^{n-1} k_1^n}{(g^n + k_1^n)^2}\frac{k_2}{k_2 + h_{p_3}}$ may take arbitrarily large (positive) values for large $V$. In this case, at least one of the eigenvalues of the Jacobian must have a positive real part, and the fixed point must therefore be unstable.

We thus conclude that, for small $V$, there is only one (stable) fixed point at the origin; we call this the **monostable regime**. As $V$ is increased, two further fixed points emerge, one stable (at high $g$), and one unstable (at low $g$); this is the **bistable regime**. The bifurcation that creates a new pair of fixed points (one stable and the other unstable) is known as a saddle-node bifurcation. Note that changes in other parameters (including $k_1, k_2, \psi$) also result in a similar bifurcation, either by stretching the $g$ nullcline or by shifting the $h$ nullcline. Therefore, in the following, we will place emphasis on variations in $V$ noting that our conclusions on the dynamics will apply equally to variations in other parameters.

#### Dynamical trajectories

Eqs. (1,2) describe an excitable system, where the magnitude of the initial perturbation away from steady-state ($g = h = 0$) determines the dynamical trajectories. In the context of the biological system, the stable fixed point at the origin ($p_1$) represents the case where the gene is unsilenced. From the stability analysis, it follows that small perturbations from this fixed point will decay rapidly and monotonically in $g$ (Figure S2a). Larger perturbations result in dynamical trajectories that follow a characteristic long path in phase space, as can be seen in Figure S2b-d. The lower branch of the nullcline, Eq. (4), is a separatrix, so that trajectories that start above it move away from the origin and towards the upper branch of the nullcline, Eq. (4). For these trajectories, it is possible for both $g, h$ to increase. The trajectories then progress along the branch, with their ultimate fate dependent on whether the system is positioned in the monostable or bistable regime. In the former (stable) regime, the trajectory "drops off the tip" and decays back towards the origin. In the latter (bistable) regime, the trajectory converges on the stable fixed point $p_2$. Examples of the range of dynamical trajectories are shown in Figure S2.

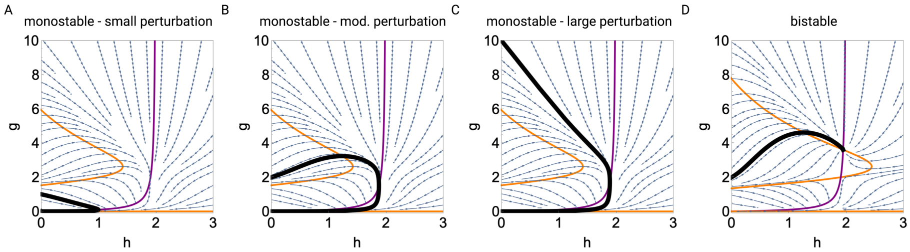


**Figure S2:** Dynamical trajectories of the rate equations (1,2) starting from an initial condition with $h = 0$ and variable $g$ in the monostable regime (panels A-C) and the bistable regime (panel D). Parameters are the same as that used in Figure S1 in the respective regimes with nullclines labelled using the same color scheme.

#### Non-zero production

The analysis thus far focused on the case where there is no production $I(t) = 0$ (only autocatalysis). More generally, the production rate may be non-zero in several scenarios. One important case is where production occurs following the presentation of a persistent dsRNA trigger, which underlies much of the experimental work on this system. Production may also be non-zero for endogenous genes, as well as for experimental systems such as multi-copy gene arrays. These systems are associated with the persistent production of dsRNAs (and thus translate to $I(t) > 0$) [1]. Persistent (albeit fluctuating) production of dsRNAs may be typical of many genes. Here, we will now study the properties of this more general situation.

When the production $I(t) > 0$ is constant $I(t) = I$, the origin $g = 0$, $h = 0$ is no longer a fixed point of the system. Instead, a stable fixed point appears at $g \approx I/\gamma_1$, $h \approx \frac{I\psi}{\gamma_2(I+\gamma_1 k_3)}$. The original nullcline at $g = 0$ shifts upwards to $g \approx I/\gamma_1$, while the second nullcline becomes stretched towards the right. If the system were positioned in the monostable regime, the increase in production rate may potentially lead to a bifurcation with the birth of a pair of stable/unstable fixed points (see Figure S3a,b). In this scenario, the system may alternate between some baseline level of gene silencing and a higher excitable permanently silenced state (similar to the dynamics observed in [3]). Finally, at even higher rates of production $I$ (where $I/\gamma_1 > k_1$), the shape of the nullcline will change and another bifurcation will occur. In this case, the system will only have a single fixed point at a high silencing level.

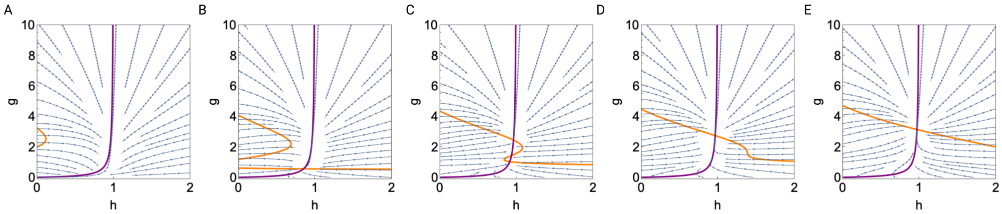


**Figure S3:** Stream plot of Eqs. (1,2) and nullclines for $\dot{g} = 0$ (orange) and $\dot{h} = 0$ (purple), setting (A) $I = 0$, (B) $I = 0.05$, (C) $I = 0.07$, (D) $I = 0.08$, and (E) $I = 0.1$. All other parameters are the same as Figure S1, with $V = 0.4$. Starting from the monostable regime (A), as $I$ increases, the lower stable fixed point moves upwards and the upper orange nullcline is stretched towards the right (B). This may result in a saddle-node bifurcation (C), and in a bistable regime similar to the one described in previous figures, but with a non-zero lower stable fixed point. As $d$ increases further, the lower stable / unstable fixed point pair may coalesce and the system will undergo another saddle-node bifurcation, with only a stable silenced fixed-point (D,E).

Going forward, we will consider the production rate $I(t)$ as a time-varying (and, occasionally, stochastic) variable. In this case, a step-like increase in $I$ (corresponding to treatment with dsRNA trigger) results in a shift of $g, h$ towards the upper-right corner of the phase space while, following cessation of dsRNA treatment, the dynamics will either return to $p_1$ (monostable regime) or shift to $p_3$ (bistable regime), as depicted in Figure S1.

#### Dynamics near the critical point

As $V$ increases, we showed above that the system transitions from a monostable to a bistable regime. This transition occurs at a specific $V = V_{\text{crit}}$ via a saddle-node bifurcation. Near the bifurcation, when $V$ is either slightly above or below $V_{\text{crit}}$, the system behaves in a special manner that will be of crucial importance for this study. Here, we will summarize some aspects of this behavior.

Consider the case where $V > V_{\text{crit}}$ and $V$ is slowly decreased until the critical point $V = V_{\text{crit}}$. Remember that the stable fixed fixed point $p_3$ is associated with positive eigenvalues $l_1, l_2 > 0$ and that the unstable fixed point $p_2$ has a negative eigenvalue $l_1 < 0$. At $V = V_{\text{crit}}$ these points coalesce to a single critical point positioned at $p_{\text{crit}}$. At $p_{\text{crit}}$, one of the eigenvalues of Eq. (7) becomes zero ($l_1 = 0$). This eigenvalue is associated with slow movement parallel to the $g$-axis. The other eigenvalue, $l_2 = -\gamma_2$, is associated with rapid convergence parallel to the $h$-axis. The dynamics of the system near the critical point are thus rendered effectively one-dimensional. To analyze the behavior in the vicinity of this point, we can consider the normal form of the saddle-node bifurcation [7]:

$$\dot{x} = x^2 - \mu \tag{10}$$

where $\mu$ is the bifurcation parameter corresponding to the scaled distance from bifurcation (Figure S4). Note that, in the case of the TI model, the parameter $\mu$ essentially captures the distance between $p_{\max}$ and the nullcline of $h$, which saturates at $h = \frac{\psi}{\gamma_2}$. Thus, applied to the current system, $\mu \propto V - V_{\text{crit}}$. We will thus consider Eq. (10) to retrieve properties of interest of the system near the critical point, which will greatly simplify the analysis. For $\mu > 0$, the normal form of the rate equation has two fixed points: a stable fixed point at $x = -\sqrt{\mu}$ and an unstable fixed point at $x = \sqrt{\mu}$. For $\mu < 0$ the system is only stable at $x = \infty$ while, for the critical case $\mu = 0$, there is a semi-stable fixed point at the origin. In the original system, $\mu > 0$ corresponds to the bistable regime, while $\mu < 0$ corresponds to the monostable regime.

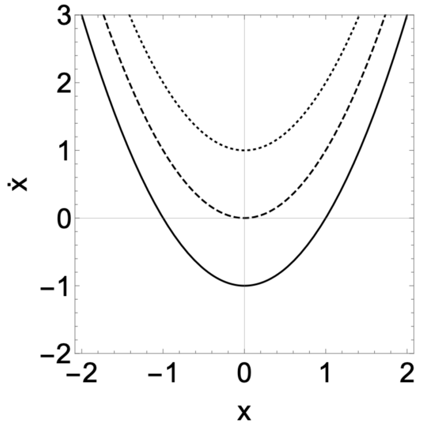


**Figure S4:** Dynamical rates of the normal form $\dot{x} = x^2 - \mu$ for different values of $\mu$ (solid: $\mu = 1$, dashed: $\mu = 0$, dotted: $\mu = -1$). When $\mu < 0$ there are no finite fixed points and the dynamics converge to $x \to \infty$ (corresponding to the monostable regime of the TI model). When $\mu > 0$ the system has a stable fixed point at $x = -\sqrt{\mu}$ and an unstable fixed point at $x = \sqrt{\mu}$ (corresponding to the bistable regime of the TI model). The case of $\mu = 0$ corresponds to the critical regime in the TI model - as $\mu \to 0$ the dynamics of the system become delayed around $x = 0$

.

A key quantity of interest when the system is in the bistable regime ($\mu > 0$) is its response to perturbations near the fixed point $x = -\sqrt{\mu}$. Setting $x' = x + \sqrt{\mu}$, we can linearize the rate equations around the fixed point to obtain:

$$\dot{x'} = -2\sqrt{\mu}x' \tag{11}$$

In response to the perturbation, the system decays back to the stable fixed point. However, as $\mu \to 0$ (that is, as $V \to V_{\text{crit}}$), the timescale of the response diverges and perturbations around the fixed point become persistent. This is the phenomenon of critical slowing down - small perturbations around the critical point become long-lived.

Crucially, for $\mu < 0$ (the monostable regime), the system may also experience a protracted delay when it transitions near $x = 0$ (a phenomenon known as a "ghost" [7]). Consider a trajectory starting at some $x_0 < 0$, which will then move towards $x \to \infty$. We can compute the delay by integrating the rate equation around the origin:

$$T = \int_0^T dt = \int_{x_0}^{\infty} \frac{dx}{x^2 - \mu} = \left. \frac{\tan^{-1}\left(\frac{x}{\sqrt{|\mu|}}\right)}{\sqrt{|\mu|}} \right|_{x_0}^{\infty} \approx \frac{\pi}{\sqrt{|\mu|}} \tag{12}$$

Thus, in the monostable regime, as $V \to V_{\text{crit}}$, the delay around the fixed point diverges as $T \sim |V - V_{\text{crit}}|^{-1/2}$. In the context of the TI model, this means that a large fluctuation from the fixed point $p_0$ induced by a trigger will follow a trajectory in phase space that incurs a long time delay $T$ close to the saddle-node bifurcation before decaying back to $p_0$.

#### Effect of noise on the dynamics

Noise in molecular reaction dynamics plays an important role in a myriad of biological processes. It may result from intrinsic fluctuations (that may occur, for example, when the number of reacting species is small) or from extrinsic fluctuations due to metabolism, stress, etc. In our case, noise is especially important since, near the critical point, its effects may become amplified. Additionally, in the deterministic (noise-free) model (1,2), the time delay $T \sim \infty$ around $V = V_{\text{crit}}$ while, in a more realistic setting, fluctuations are likely to result in $T$ remaining finite. To address the role of fluctuations, we therefore modified Eqs. (1,2) to include noise terms:

$$dg = \left( I(t) + V \frac{g^n}{k_1^n + g^n} \frac{k_2}{k_2 + h} - \gamma_1 g \right) dt + \sigma_1 g \, dW_1 \tag{13}$$

$$dh = \left( \psi \frac{g}{k_3 + g} - \gamma_2 h \right) dt + \sigma_2 h \, dW_2 \tag{14}$$

where $\sigma_1$, $\sigma_2$ are the respective noise amplitudes, and $W_1$, $W_2$ denote (uncorrelated) one-dimensional Wiener processes [8]. Here we consider multiplicative noise, which is common in biological settings and has the attractive property that it cannot result in negative concentrations. We note that considering additive noise would lead to identical conclusions. Near the critical point, there is little variation in $(g, h)$. Therefore, to study the impact of noise on the dynamics, we may transform Eq. (10) to the corresponding stochastic differential equation:

$$dx = (x^2 - \mu) dt + \sigma W \tag{15}$$

where $W$ is a one-dimensional Wiener process and $\sigma$ denotes the effective noise amplitude. In the bistable regime, $T$ is governed by the escape rate from the stable fixed point at $x = -\sqrt{\mu}$ to the stable fixed point at infinity.

Here, we may make an analogy to the thermally-assisted escape rate of an over-damped particle confined to a well with the potential

$$\Phi = -\frac{x^3}{3} + \mu x \tag{16}$$

The delay time $T$ can be estimated by using Kramer's approximation for the escape rate from a potential well:

$$T \approx \frac{2\pi}{\sqrt{\ddot{\Phi}(-\sqrt{\mu}) |\ddot{\Phi}(\sqrt{\mu})|}} e^{2(\Phi(\sqrt{\mu}) - \Phi(-\sqrt{\mu}))/\sigma^2} = \frac{\pi}{\sqrt{\mu}} \exp\left[ \frac{8}{3\sigma^2} \mu^{3/2} \right] \tag{17}$$

Thus, as the amplitude increases beyond the bifurcation point $V > V_{\text{crit}}$, there is a steep rise in the escape time $T$, increasing as $\ln T \sim \mu^{3/2}$ (and thus $\ln T \sim (V - V_{\text{crit}})^{3/2}$). The approximations for scaling behavior in the bistable regime ($\mu > 0$, where $\ln T \sim \mu^{3/2}$ in the noisy case) and monostable regime ($\mu < 0$, where $T \sim |\mu|^{-1/2}$ in the deterministic case) can be generalized to any $\mu$ under the presence of noise, as shown recently by Hathcock and Sethna [2]. In this case, $T$ scales as

$$T \approx 2^{1/3} \pi^2 \sigma^{-2/3} \left( \text{Ai}^2 [2^{2/3} \frac{\mu}{\sigma^{4/3}}] + \text{Bi}^2 [2^{2/3} \frac{\mu}{\sigma^{4/3}}] \right) \tag{18}$$

where $\text{Ai}, \text{Bi}$ denote the first and second Airy functions. The behavior of Eq. (18) resembles the Kramers escape time for the bistable regime and deterministic scaling for the monostable regime, and can be used to study the escape rate for arbitrary $\mu$ in the vicinity the critical point, regardless of the specific model details.

The incorporation of noise into the dynamical rate equations allows us to consider other statistics of interest, including the variance of sample trajectories around the stable fixed-point in the bistable regime, and the autocorrelation of individual trajectories (defined as the correlation between the values of sample paths over a fixed time-lag, Figure S5). Both quantities diverge near the critical point, and there is extensive literature on the nature of this divergence (as these are considered as "early indicators" for critical transitions) [5]. In particular, for a saddle-node bifurcation, the variance of sample trajectories around the critical for Eq. (10) diverges as $|V - V_{\text{crit}}|^{-1}$.

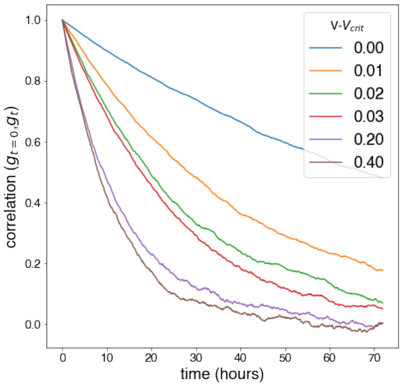


**Figure S5:** Autocorrelation of individual sample paths near the critical point during a single generation, three generations after initiation of silencing, for different values of $V - V_{\rm crit}$, measured by the Pearson correlation between $g(t=0)$ and $g(t = \text{lag})$. In order to prevent trajectories from crossing the critical point, noise was reduced to $\sigma = 0.0001$. Other parameters and simulation details are identical to Figure S2, and the generation time $\tau$ is provided in Table 3.

#### 2 Toggle-Inhibitor-Competition (TIC) model

Close to the saddle-node bifurcation, the TI model provides a framework to describe much of the observed phenomenology of gene silencing. However, as discussed in the main text, its application to the experimental system requires a high degree of fine-tuning, which is hard to motivate. Here, we develop a modified framework based on competition for silencing machinery, in which the system self-organizes around the saddle-node bifurcation without fine-tuning.

In the Toggle-Inhibitor model, emphasis was placed on the silencing of a single gene. The Toggle-Inhibitor-Competition (TIC) model considers the dynamics of all genes, accounting for their competition over shared synthesis resources. Let $i = 1, ..., N$ denote the indices of genes in the ensemble of endogenous and exogenous genes that may become silenced, with the corresponding concentrations of gene-specific siRNAs and silencing modification marks denoted by $g_i, h_i$. The collective dynamics are then given by the coupled set of rate equations

$$\dot{g_i} = I_i + \frac{V_{\rm tot}}{C(\{g_i, h_i\})} \frac{g_i^n}{k_1^n + g_i^n} \frac{k_2}{k_2 + h_i} - \gamma_1 g_i \tag{19}$$

$$\dot{h_i} = \psi \frac{g_i}{k_3 + g_i} - \gamma_2 h_i \tag{20}$$

where the "cost" function $C$ is proportional to the overall abundance of siRNAs,

$$C = \sum_{i=1}^{N} c(g_i, h_i) = \sum_{i=1}^{N} g_i \tag{21}$$

Here, for simplicity, we have taken the model parameters to be gene-independent, turning later to consider the effect of variations. As we will see below, the precise form of $C$ is not important for the conclusions. It is only important that $C$ increases with the number of actively silenced genes. Moreover, alternative models where other parameters are modulated, such as a model where $k_1$ increases with $C$, behave in an almost identical manner for the considered properties of interest.

In this model, we consider the ensemble of production rates $I_i$ as a random variable, where stochasticity may arise due to fluctuating environmental conditions, or internal stochastic events such as transcriptional bursting. To simplify the analysis, we will consider a random process where, at a given generation, $I_i = 0$ with probability $1 - q$ or $I_i = L$ (for a fixed $L > 0$) with probability

$q$. Further, with silencing events considered as infrequent, we may assume that $q \ll 1$. We also assume that silencing events occur only when the gene is unsilenced; that is, when $g_i, h_i$ are near the origin. We note that, for large $N$, fluctuations in $C$ due to such stochastic dynamics will be small.

Finally, similar to Eqs. (13,14), in the presence of noise, the corresponding equations take the form:

$$dg_i = \left( I_i + \frac{V_{tot}}{C(\{g_i, h_i\})} \frac{g_i^n}{k_1^n + g_i^n} \frac{k_2}{k_2 + h_i} - \gamma_1 g_i \right) dt + \sigma_1 g_i \, dW_{i_1} \tag{22}$$

$$dh_i = \left( \psi \frac{g_i}{k_3 + g_i} - \gamma_2 h_i \right) dt + \sigma_2 h_i \, dW_{i_2} \tag{23}$$

#### Global dynamics and steady state

To analyze the stability of Eqs. (19,20), we will consider a system starting from either (i) very few silenced genes (and therefore a very small cost $C = C_{\text{min}}$), or (ii) many silenced genes (with very large $C = C_{\text{max}}$). As before, we denote by $V_{\text{crit}}$ the (this time per-gene) synthesis rate at which the system transitions from monostability to bistability. In case (i), the effective synthesis rate $\frac{V_{\text{tot}}}{C_{min}}$ may be very large, placing the system in the regime of bistability ($\frac{V_{\text{tot}}}{C_{\text{min}}} > V_{\text{crit}}$). Thus, new stochastic silencing events ($I_i = L$) will result in stable silencing, driving an increase in the cost function ($\dot{C} > 0$). On the other hand, in case (ii), the system is in the monostable regime ($\frac{V_{\text{tot}}}{C_{\text{max}}} < V_{\text{crit}}$), where genes gradually become silenced and the cost function will fall ($\dot{C} < 0$). Thus, over time, the system will converge towards a dynamic steady-state (with $\dot{C} = 0$) at some intermediate value of the cost function $C = C_{\text{st}}$.

What is the steady-state value $C_{\text{st}}$? We will now show that, for a wide range of parameter values, $C_{\text{st}} \approx C_{\text{crit}}$, where

$$\frac{V_{\text{tot}}}{C_{\text{crit}}} = V_{\text{crit}} \tag{24}$$

To see why this is the case, suppose that the (average) number of silenced genes at steady-state is given by $M < N$. Let us denote by $\lambda$ the arrival rate of new silencing events, where according to the model, $\lambda = q(N - M)$. At steady-state the system must adhere to Little's law [4]:

$$M = \lambda T = q(N - M)T \tag{25}$$

where $T$ is the average silencing duration of an individual gene. In other words, the flux of new silenced genes must equate to the number that become de-silenced. When the system is near the critical point,

$$C_{\text{crit}} \approx M c_{\text{crit}} \tag{26}$$

where $c_{crit} = c(p_{\text{crit}})$ is the cost for an individual gene at the stable fixed point (here $c(p_{\text{crit}}) = g_{\text{crit}}$). Combining Eqs. (24,25,26), we obtain an (approximate) equation for T as

$$T = \frac{1}{q\left(\frac{N c_{\text{crit}} V_{\text{crit}}}{V_{\text{tot}}} - 1\right)} \tag{27}$$

To gain some intuition for Eq. (27), consider the case where $N = 1000$ genes and $q = 0.01$ per generation, so that the arrival rate of new silencing events, when no genes are currently silenced, is 10 genes per generation. Setting $c_{\text{crit}} = 1$, $V_{\text{crit}} = 1$ (note that these depend on circuit parameters), we obtain the equation, $T = \frac{100 \times V_{\text{tot}}}{1000 - V_{\text{tot}}}$. For $V_{\text{tot}} > 10$ and $V_{\text{tot}} \ll 1000$, the right-hand side increases linearly with $V_{\text{tot}}$ and is on the order of several (to several dozens of) generations. This equation will be satisfied near the critical point. The reason for this is that this timescale is slower than the typical timescale in the monostable regime, yet it will be realized in the vicinity of the critical point due to the rapid increase in $T$ in this region (see Eq. (17)). More generally, taking the limit $N \to \infty$ and $q \to 0$ (with $\lambda = qN$ constant),

$$T = \frac{V_{\text{tot}}}{\lambda c_{\text{crit}} V_{\text{crit}}} \tag{28}$$

In this limit, any $V_{\text{tot}} > \lambda c_{\text{crit}} V_{\text{crit}}$ will result in a transgenerational timescale, which would lead to the system converging to a steady-state positioned near the critical point. Finally, from Eqs. (24,26), it follows that

$$M = \frac{V_{\text{tot}}}{c_{\text{crit}} V_{\text{crit}}} \tag{29}$$

so that the average number of silenced genes at steady-state increases linearly with $V_{\text{tot}}$.

It is important to note that there were certain assumptions made in the above derivations. In particular, we assumed a large separation of timescales between that of the steady-state, Eq. (27), and the rapid turnover timescale in Eqs. (1,2) and the generalization (22,23). We also assumed that the system is in the monostable regime when $C = C_{\text{max}}$ and bistable when $C = C_{\text{min}}$. This might not be the case if, for example, there is a large decrease in $N$ (as may occur, for example, when piRNAs are deleted). In this case, the system may settle in the bistable state, as can be seen in Figure S6.

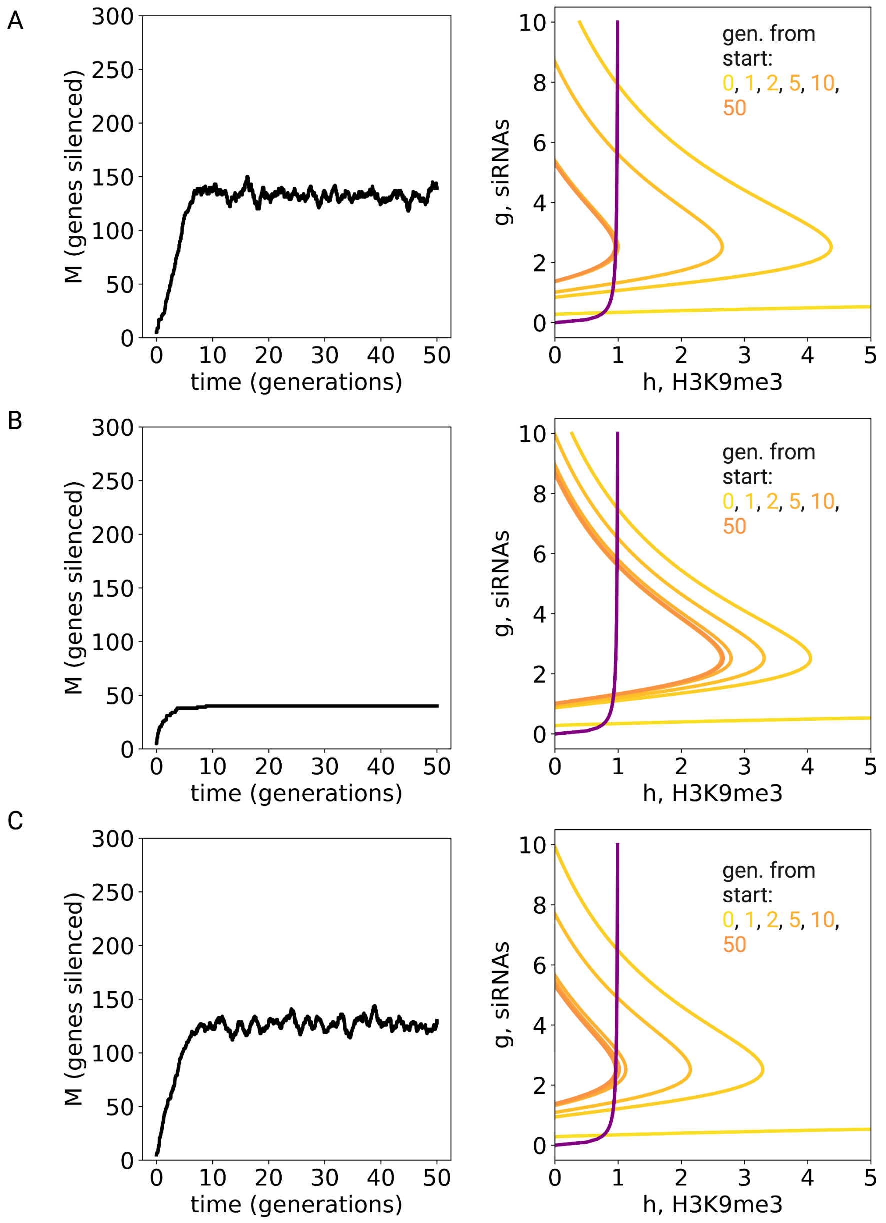


**Figure S6:** Stochastic simulations of the TIC model (as in Figure 4A,B), taking either (A) an infinite pool of genes that can be silenced, or (B) a finite pool of magnitude $N = 40$ or (C) $N = 400$. When all candidate genes become silenced (as in panel B), the system may stabilize in the bistable region.

#### Variation in parameters

Finally, we turn to consider the role of potential variation in the model parameters. Consider a system that is organized around some steady-state cost $C = C_{\text{st}}$. So far, we assumed that all genes have identical parameters, and that $C$ is constant at steady-state. However, in a realistic setting, this assumption may break down. One possibility is that there is variation in biochemical parameters between genes, which may be static over time. Another possibility is that there are global fluctuations, such as fluctuations in $V_{\text{tot}}$, which can lead to variation in $C_{\text{st}}$ over time and possibly between worms and worm lineages. Another source of variation can be changes in $C_{\text{st}}$ due to variation in the arrival of silencing memories. This variation can affect both the steady-state cost $C_{\text{st}}$, as well as the silencing duration of individual genes.

For global and stochastic fluctuations in parameters values, we would still expect that the system will settle near the critical point due to the same argumentation as in previous sections, even in the presence of variation in parameters and between genes. Settling near the bifurcation is a robust property that holds for a wide range of parameter values, and so will be unaffected by fluctuations in global parameter values. For variation in individual genes, we should consider that Little's law must hold for the genes that are transgenerationally silenced (as they contribute to the cost $C$). As an illustrative example, we can consider a population of genes where half of the genes have a much smaller value of $V_{\mathrm{tot}}$ than the other genes. In this case, at steady-state, the genes with the higher $V_{\mathrm{tot}}$ will satisfy Eq. (28) (with an arrival rate of $\lambda/2$), while the genes with the lower $V_{\mathrm{tot}}$ will be in the monostable regime and thus will contribute only little to $C$. The system will thus settle near the critical point for the genes that have stable transgenerational inheritance.

To extend this argument, we can ask how continuous variation in model parameters affects the global steady-state of the system (Figure S7). To this end, we considered the situation in which $V_{\mathrm{tot}}$ is Gaussian distributed in the population of genes, with an increasing standard deviation $b$. As can be seen in Figure S7, as $b$ increases, the value of $\frac{V_{\mathrm{tot}}}{C}$ decreases slightly below the critical value, and the value of $\frac{V_{\mathrm{tot}}}{C}$ of the silenced genes remains near the critical $V_{\mathrm{crit}}$. Thus, variation between genes leads to a subgroup of genes closer to the monostable regime (which cannot be transgenerationally silenced), and another subgroup that has longer transgenerational silencing. Similar results hold for variation in all model parameters. Tuning near the critical point is thus robust to variation in parameters between genes.

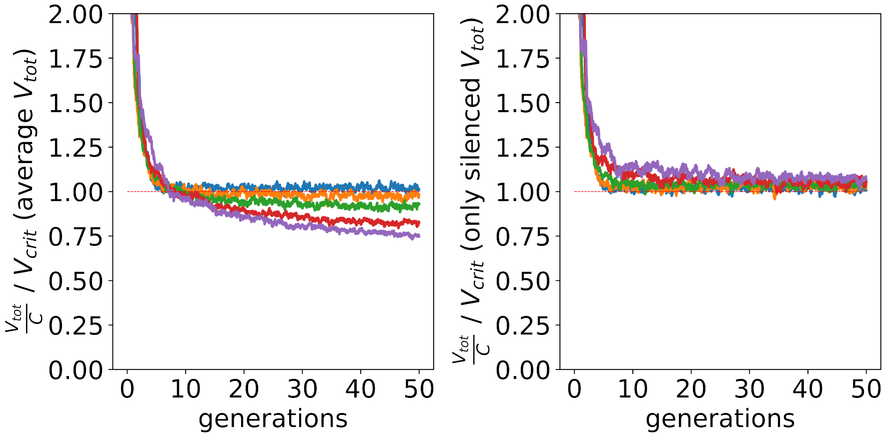


**Figure S7:** (A) Change in $\frac{V_{\mathrm{tot}}}{C}$ as a function of generation number in the situation where memory population dynamics were simulated as in Figure 4 in the main text, but where $V_{\mathrm{tot}}$ for individual genes is drawn from a normally-distributed random variable with $0\%, 5\%, 10\%, 20\%, 30\%$ standard deviation around the mean (blue, orange, green, red, and purple line respectively). (B) In all cases, the value of $\frac{V_{\mathrm{tot}}}{C}$ of the silenced genes remains close to $V_{\mathrm{crit}}$.

We next considered the effect of variation in model parameters on variation in silencing duration. Here, once again, one may take advantage of the normal form of the dynamics of the system near the saddle-node bifurcation, Eqs. (10,15). We can capture the variation by considering $\mu$ in the normal form of the model as a random variable, drawn from some distribution $F_\mu(\mu)$. To derive analytical results, we can divide the contributions from where $F_\mu$ is in the monostable regime (where $\mu < 0$), denoted $F_{\mu_-}$, and from where $F_\mu$ in the bistable regime (where $\mu > 0$), denoted $F_{\mu_+}$. Starting with the contribution from the former, in this case $T \approx a|\mu|^{-1/2}$ (where $a$ is a constant), from which it follows that $\mu = -\frac{a^2}{T^2}$. It therefore follows that the probability density for $T$ is given by

$$\Pr(T) = F_{\mu_-}\left(-\frac{a^2}{T^2}\right)\left|\frac{d\mu}{dT}\right| = F_{\mu_-}\left(-\frac{a^2}{T^2}\right)\frac{2a^2}{T^3} \approx F_{\mu_-}(0)\frac{2a^2}{T^3} \tag{30}$$

where the last approximation is relevant for large $T$. Thus, the part of the distribution where $\mu < 0$ contributes a delay distribution that scales as $T^{-3}$.

When variations place the system in the bistable regime, with the distribution $F_{\mu+}$, it follows from the analysis above that $T \approx A\mu^{-1/2}e^{B\mu^{3/2}}$ (with constants $A$, $B$). In this case, the tail of the distribution is dominated by large $\mu$. Therefore, to simplify the analytical expressions, we will neglect the $\mu^{-1/2}$ prefactor and simply take $T \approx Ae^{B\mu^{3/2}}$, or $\mu \approx \left(B^{-1}\ln(T/A)\right)^{2/3}$. In this case, the probability density is given by

$$\Pr(T) \approx F_{\mu+}\left(\left(\frac{\ln(T/A)}{B}\right)^{2/3}\right)\left|\frac{d\mu}{dT}\right| \approx F_{\mu+}\left(\left(\frac{\ln(T/A)}{B}\right)^{2/3}\right)\frac{2}{3BT}\frac{1}{\sqrt[3]{\frac{\ln(T/A)}{B}}} \tag{31}$$

The term $\sqrt[3]{\frac{\ln(T/A)}{B}}$ varies slowly at large $T$ compared with the $1/T$ dependence. The term $F_{\mu+}\left(\left(\frac{\ln(T/A)}{B}\right)^{2/3}\right)$ depends on the form of the distribution. However, typically, it will decay very slowly when the variance of the distribution is large. For example, in the case of a normal distribution, $F_\mu = \frac{e^{-(\mu/b)^2/2}}{\sqrt{2\pi}b}$, the survival function is then given by

$$\mathrm{S}(T) = \frac{G(T)}{G(T_{\min})} \tag{32}$$

where:

$$G(T) = \frac{\left(\frac{\ln(T/A)}{B}\right)^{2/3} E_{\frac{1}{2}}\left(\frac{1}{2b^2}\left(\frac{\ln(T/A)}{B}\right)^{4/3}\right)}{4\sqrt{2\pi}b} \tag{33}$$

As the standard deviation $b$ increases, this function becomes effectively constant for large $T$. (Note that other plausible distributions, such as log-normal, show similar behavior.) In this case, the tail of distribution $\Pr(T)$ becomes entirely dominated by the $1/T$ dependence, resulting in a heavy tail with a nearly-flat survival function.

Finally, even for a fixed parameter set, there would be variation due to the noise terms in Eqs. (13,14). However, this variation does not result in long tails (see Ref. [2] for derivations of the distributions, including comparison with stochastic simulations).

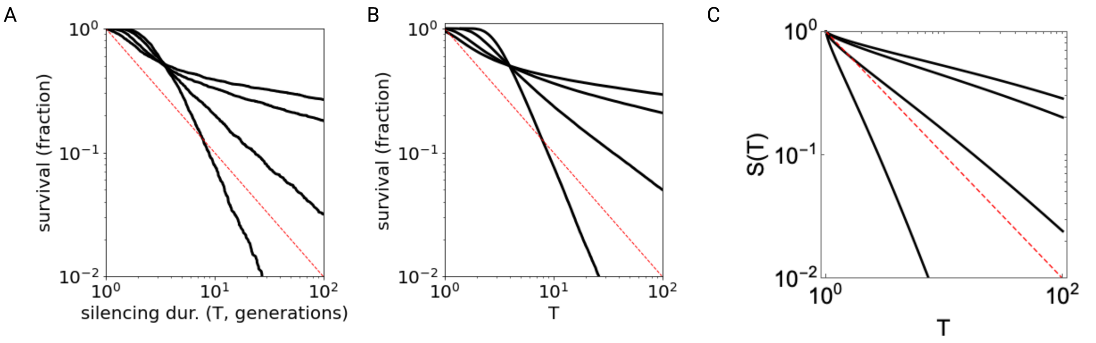


**Figure S8:** (A) Survival time distribution obtained from stochastic simulation of Eqs. (13,14), taking the parameters as in Figure S2, but setting $V \sim \text{Normal}(V_{\text{crit}}, b)$, with $b = 0.01, 0.02, 0.04, 0.06$ (lower to upper lines). Noise is set as in other simulations to $\sigma_1 = \sigma_2 = 0.03$. Trajectories start at $(g_0, h_0) = (3, 0)$ and are considered silenced until the silencing metric $s(g, h)$ drops below the threshold $s_m$. This results in a distribution of survival times, which is depicted on a log-log plot. (B) The survival distribution of $T$ calculated from Eq. (18), taking $\sigma = 2$ and drawing $\mu$ from a normal distribution $\mu \sim \text{Normal}(0, b)$ with $b = 1, 2, 4, 6$. (C) Predicted survival time distribution $\mathrm{S}(T)$ as defined by Eqs. (32,33), setting $b = 0.01, 0.02, 0.04, 0.06$ (lower to to upper lines). Note that this approximation, which captures the bistable portion of $V$, is appropriate only for the slope of tails of the survival distributions depicted in panel A. Here, we have set $A = 1$, $B = 482$. Adjusting $A$ does not sensitively affect the outcome as long as $\ln T \gg \ln A$, while $B$ is inferred from simulations of average silencing duration vs. $V - V_{\text{crit}}$. The dashed red line denotes $t^{-1}$. We note that, as the variation $b$ increases, the tail of the survival distribution asymptotes towards a flat line parallel to the time axis.

#### More complex model that incorporates pUGylation dynamics

While our analysis focused on the simple two dimensional TI model (provided by Eqs. 1,2), our conclusions regarding self-tuning near the critical point apply to a much wider class of models. The only necessary ingredient is the competition over autocatalysis resources between the (excitable) silencing memories, which tune a control parameter for a saddle-node bifurcation. The dynamics can be given by a complex and high-dimensional interaction, rather than by the simple two dimensional system provided by the TI model. To demonstrate this, we show that our conclusions are effectively unchanged when we consider a more complex model for silencing dynamics, which incorporates the important process of pUGylation of template mrNAs. It has been demonstrated that siRNAs are produced from template mRNAs which have been tagged with poly(UG) tails [6]. The tagging itself is performed in a manner which is directed by existing siRNAS [6]; it is thus an autocatalytic process. Additionally, we assume that the production of template mRNAs is inhibited by the silencing chromatin marks. Based on these interactions, we propose the following rate equations:

$$dg_i = \left( I_i + \frac{V_{tot}}{\sum g_i} y_i - \gamma_1 g_i \right) dt + \sigma_1 g_i \, dW_{i_1} \tag{34}$$

$$dh_i = \left( \frac{g_i}{g_i + k_3} - \gamma_2 h_i \right) dt + \sigma_2 h_i \, dW_{i_2} \tag{35}$$

$$du_i = \left( \frac{k_2}{h_i + k_2} - \gamma_3 u_i \right) dt + \sigma_3 u_i \, dW_{i_3} \tag{36}$$

$$dy_i = \left( \frac{(g_i u_i)^n}{k_1^n + (g_i u_i)^n} - \gamma_4 y_i \right) dt + \sigma_4 y_i \, dW_{i_4} \tag{37}$$

where $g_i, h_i, u_i, y_i$ correspond to siRNA concentration, concentration of chromatin silencing marks, mRNA levels, and mRNA molecules tagged by pUG tails for gene $i$ ($i \in 1, ..., N$). As before, we simulated the model with stochastic activation effects. Despite the added complexity, this model also self-tunes near the saddle-node bifurcation point (Fig. S9).

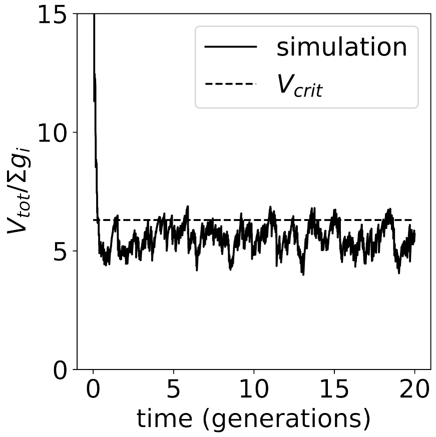


**Figure S9:** Simulation of Eqs. 34,35,36,37, taking all parameters as in Tables 1,2,3, with the simulation starting with no silenced genes. The effective $V_{tot}/\sum g_i$ converges rapidly to the vicinity of the critical value.

#### Memory system timescales

The model has two important timescales. The first timescale is the fast timescale of dilution/turnover of memory components, given by the decay times $\gamma_1^{-1}, \gamma_2^{-1}$. The second timescale is the slow memory duration timescale $T$. These timescales govern dynamical properties of the memory system. Consider a step change in a (global) model parameter, such as the overall synthesis capacity $V_{\text{tot}}$, which will result in a change in the steady-state memory pool size $M$. If this is a down-step (and $M$ decreases), the system will be transiently in the monostable regime, and genes will be removed according to the fast dilution timescale (Figure S10). On the other hand, an up-step in $V_{\text{tot}}$ will result in the system transiently moving to the bistable regime. Genes will then re-accumulate according to the stochastic arrival rate $\lambda$, with a timescale given by $\frac{M}{\lambda} = T$. Measuring these timescales may be possible in experiments where silencing is known to be globally perturbed, such as in conditions of stress.

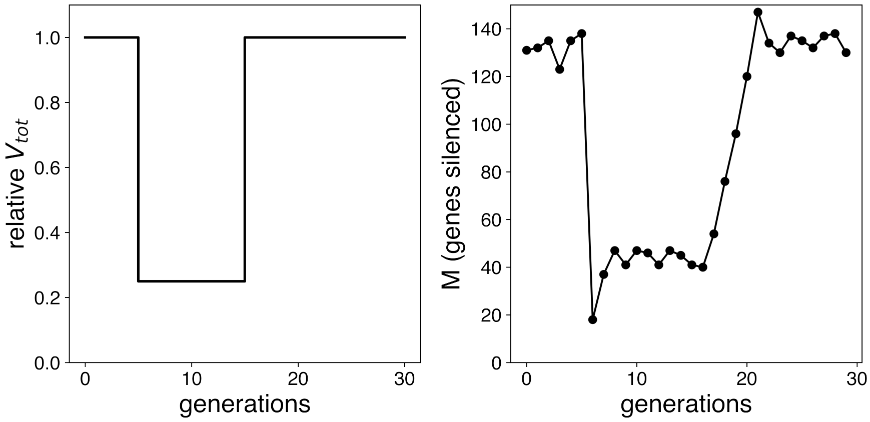


**Figure S10:** Step changes in parameters result in asymmetric silencing dynamics in the TIC model. A down-change in the silencing pool size $M$ (such as due to a down-step in $V_{\text{tot}}$) is driven by a transition of the system to the monostable regime and thus has a fast timescale. An up-change in $M$, on the other hand, requires re-accumulation of silenced genes and thus has a longer timescale, on the order of $T$. This is evident by comparing the effect of down- and up-steps in $V$ (left) on the silencing pool size $M$ (right). Simulation parameters are provided in Tables 1, 2, and 3.

#### Arrival-rate is independent of memory duration

A final intriguing possibility is that the dependence on memory arrival rate $\lambda$ in Eq. (28) can be completely removed by adding a feed-forward interaction between the arrival of a new memory event and synthesis capacity $V_{\text{tot}}$, such that $V_{\text{tot}} = \eta\lambda$ for some proportionality constant $\eta$. This may occur, for example, if external triggers or internal activation pulses also encourage the production of more autocatalysis enzymes. In this case, Eq. (28) becomes:

$$T = \frac{\eta}{\lambda c_{\text{crit}} V_{\text{crit}}} \tag{38}$$

which depends only on internal biochemical parameters.

| Parameter | Value | Units | Comment |
|---|---|---|---|
| $V$ | 0.45, 0.55, 0.8 | $[\mathrm{AU}]\mathrm{hr}^{-1}$ | monostable, critical, bistable |
| $n$ | 3 | unitless | |
| $k_1$ | 2 | [AU] | |
| $k_2$ | 2 | [AU] | |
| $k_3$ | 0.1 | [AU] | |
| $\psi$ | 1 | $[\mathrm{AU}]\mathrm{hr}^{-1}$ | |
| $\gamma_1$ | 0.1 | $\mathrm{hr}^{-1}$ | dilution timescale |
| $\gamma_2$ | 0.1 | $\mathrm{hr}^{-1}$ | dilution timescale |
| $\gamma_3$ | 0.1 | $\mathrm{hr}^{-1}$ | dilution timescale |
| $\gamma_4$ | 0.1 | $\mathrm{hr}^{-1}$ | dilution timescale |
| $I$ | 0.3 | $[\mathrm{AU}]\mathrm{hr}^{-1}$ | dsRNA trigger |
| $I$ | 0 | $[\mathrm{AU}]\mathrm{hr}^{-1}$ | no dsRNA trigger |
| $\sigma_1$ | 0.03 | $\mathrm{hr}^{-1/2}$ | |
| $\sigma_2$ | 0.03 | $\mathrm{hr}^{-1/2}$ | |
| $\sigma_3$ | 0.03 | $\mathrm{hr}^{-1/2}$ | |
| $\sigma_4$ | 0.03 | $\mathrm{hr}^{-1/2}$ | |

**Table 1:** Simulation parameters for the TI model.

### 3 Details of numerical simulations

To simulate model equations, we used either the Euler method (for deterministic ODEs) or the Euler-Maruyama method (for SDEs), taking $dt = 0.1\mathrm{hr}$. All other parameters are provided in Tables 1-3. Silencing strength was estimated by taking $s = \sqrt{g^2 + h^2}$, and a gene was considered silenced if $s > s_{\mathrm{silenced}}$. For selection experiments, a gene would be considered to be strongly silenced if $s > s_{\mathrm{select}}$. Parameter values are provided in Table 3.

#### Simulations of directed vs. random selection

To simulate experimental selection (as in Figure 3) we considered the setting where, in each generation, $N = 250$ individual worms are simulated. In order to simulate the transfer of random offspring to new plates, at the end of the generation, another $N = 250$ worms were sampled with replacements, either from the entire population (random selection) or from worms with strong silencing (directed selection).

#### Simulations of TIC model

The TIC model with stochastic arrivals was simulated by taking a fixed mean arrival rate of new silencing events, given by $\lambda$, resulting in a Poisson distribution of silencing events per generation. Silencing events were simulated by adding silenced genes with $g = 3, h = 0$. Silenced genes were removed from the simulation once they became de-silenced.

#### Simulation parameters

Simulation parameters (including amplification rates and noise magnitudes) are specified in Tables 1-3. Note that the important phenomena associated with the model - namely self-tuning near the critical point - is a generic property that holds for a wide range of parameters and noise levels.

| Parameter | Value | Units | Comment |
|---|---|---|---|
| $V_{\text{tot}}$ | 200 | $[\text{AU}]\text{hr}^{-1}$ | |
| $\lambda$ | 20 | events generation$^{-1}$ | |

**Table 2:** Additional simulation parameters for the TIC model

| Parameter | Value | Units | Comment |
|---|---|---|---|
| generation length | 72 | hr | |
| $s_{\text{silenced}}$ | 0.5 | [AU] | threshold on $s$ for whether gene is silenced |
| $s_{\text{select}}$ | 3 | [AU] | threshold on $s$ for directed selection |

**Table 3:** Other simulation parameters.

### References


[1] Cristina Cruz and Jonathan Houseley. "Endogenous RNA interference is driven by copy number". In: *Elife* 3 (Feb. 2014), e01581.

[2] David Hathcock and James P Sethna. "Reaction rates and the noisy saddle-node bifurcation: Renormalization group for barrier crossing". In: *Phys. Rev. Research* 3.1 (Feb. 2021), p. 013156.

[3] Adam Klosin et al. "Transgenerational transmission of environmental information in C. elegans". en. In: *Science* 356.6335 (Apr. 2017), pp. 320–323.

[4] J D C Little. "A Proof for the Queuing Formula: L = $\lambda$W". In: *Oper. Res.* (1961).

[5] Marten Scheffer et al. "Early-warning signals for critical transitions". en. In: *Nature* 461.7260 (Sept. 2009), pp. 53–59.

[6] Aditi Shukla et al. "poly(UG)-tailed RNAs in genome protection and epigenetic inheritance". en. In: *Nature* 582.7811 (June 2020), pp. 283–288.

[7] Steven H Strogatz. *Nonlinear Dynamics and Chaos with Student Solutions Manual: With Applications to Physics, Biology, Chemistry, and Engineering, Second Edition.* en. CRC Press, Sept. 2018.

[8] N G Van Kampen. *Stochastic Processes in Physics and Chemistry.* en. Elsevier, Nov. 1992.
